# CUT&Tag for high-resolution epigenomic profiling from a low amount of *Arabidopsis* tissue

**DOI:** 10.1101/2024.07.29.604300

**Authors:** Yixuan Fu, Marc W. Schmid, Sara Simonini

**Author notes:** Author for correspondence: Sara Simonini.

## Abstract

**Background:** The genome-wide profiling of chromatin states that are defined by different histone post-translational modifications, known as epigenomic profiling, is crucial for understanding the epigenetic regulations of gene expression, both in animal and plant systems. CUT&Tag (Cleavage Under Targets and Tagmentation, [1]) is a novel enzyme-tethering method for epigenomic profiling, initially developed for mammalian cells. CUT&Tag has several advantages compared to the most commonly used epigenomic profiling methods such as Chromatin Immunoprecipitation followed by high-throughput sequencing (ChIP-seq). CUT&Tag allows epigenenomic profiling from a much less amount of starting material compared to ChIP-seq. CUT&Tag is based on the *in situ* cleavage of DNA by enzymes tethered to antibodies, while in ChIP-seq, the cleavage is done by a nearly random fragmentation step. In theory, this difference in the way of cleaving DNA allows CUT&Tag to reach a higher resolution compared to ChIP-seq. Therefore, CUT&Tag holds the potential to profile the genome-wide distribution at a high resolution even from a small amount of plant tissues.

**Results:** We profiled the genome-wide distribution of three histone modifications, H3K27me3, H3K4me3 and H3K27Ac, from a few seedlings of *Arabidopsis* that weighed around 0.01 grams. By comparing the H3K27me3 profiles generated from ChIP-seq and CUT&Tag, we showed that CUT&Tag and ChIP-seq capture the same broad lines of the epigenomes, but they also revealed different sets of peaks. Analysis using the CUT&Tag datasets for the three histone modifications revealed their genomic locations and their relationship with the gene expression level, which are consistent with the expected effect of these histone marks on gene transcription. By comparing to the nucleosome occupancy data, we show that CUT&Tag reached nucleosomal resolution, a much higher resolution than ChIP-seq. In the end, we presented that the increased resolution of CUT&Tag could better reveal the exon enrichment of histone modifications and the epigenetic states of the +1 nucleosome, showing benefits and advantages that this technique could bring to the field of plant epigenetics and chromatin study in general.

**Conclusion:** CUT&Tag is a valid, easy-to-perform, cost-effective, and reliable approach for efficient epigenomic profiling in *Arabidopsis*, even with limited amount of starting material and provides a higher resolution compared to ChIP-seq. Because the CUT&Tag protocol starting input is isolated nuclei, it is also applicable to other model and non-model plants.

## Background

The profiling of the genome-wide occupancy of histone post-translational modifications and the binding of transcription factors is crucial for understanding chromatin and epigenetic regulation in multicellular organisms. The most commonly used method for chromatin profiling is Chromatin Immunoprecipitation followed by high-throughput sequencing (ChIP-Seq; [2,3]). In this approach, DNA-protein interactions are preserved by crosslinking with formaldehyde, then the nuclei are extracted, chromatin is fragmented by sonication, and an antibody that targets the epitope of interest is added. Subsequently, the antibody-bound chromatin is recovered, and DNA is purified, adapter-ligated, amplified to generate a library, and sequenced through Next Generation Sequencing. A major limitation in using this technique in plant cells is the requirement of grams of input material. This limit brings obstacles to the studies where only a small amount of input material is available, for example, the studies of the variation of epigenetic states between individual plants and the studies of epigenetic states in specific tissues that are small and hard to collect, like embryos and meristems.

People have developed several improved versions of ChIP-seq to address its material limitations. These include ULI-ChIP [4] and ChIPmentation [5]. Among these improved versions of ChIP-seq, ChIPmentation has gained some popularity and has been successfully applied to plant systems [5,6] ChIPmentation uses the same workflow as ChIP-seq, but after the immunoprecipitation step, a Tn5 transposase is added to add adaptors to the ends of the DNA around the targeted nucleosomes. Primers binding to the adaptors are then used to amplify the tagmented fragments through PCR reactions. ChIPmentation has a reduced material requirement to a magnitude of 10^5 cells. However, these improved versions of ChIP-seq still follow the same strategy as the original ChIP-seq and thus are similarly affected by technical issues, such as the limit of the fragmentation step, which causes material loss and a genome-wide background [7]. A recently developed method based on the enzyme-tethering strategy, CUT&Tag (short for Cleavage Under Targets and Tagmentation) [1], provides another valid alternative solution for chromatin profiling. CUT&Tag uses unfixed or lightly fixed cells or nuclei as input and a primary antibody to recognize the epitopes of interest - for example, histone post-translational modifications-, followed by a secondary antibody for signal amplification. Subsequently, a hyperactive Tn5 transposase-Protein A (pA-Tn5) fusion protein loaded with DNA adaptors, is added. The protein A part of pA-Tn5 binds to antibodies with high affinity, thus tethering the pA-Tn5 enzyme to the epitope of interest. After activation by magnesium, the hyperactive Tn5 transposase cleaves local genomic DNA and adds the DNA adapters to the cleaved ends in a process called tagmentation. After DNA extraction, the tagmented and adapter-ligated DNA is amplified by PCR with full-length oligos containing the entire sequencing adaptors, which results in sequencing-ready libraries (Fig. 1).

**Figure 1.**
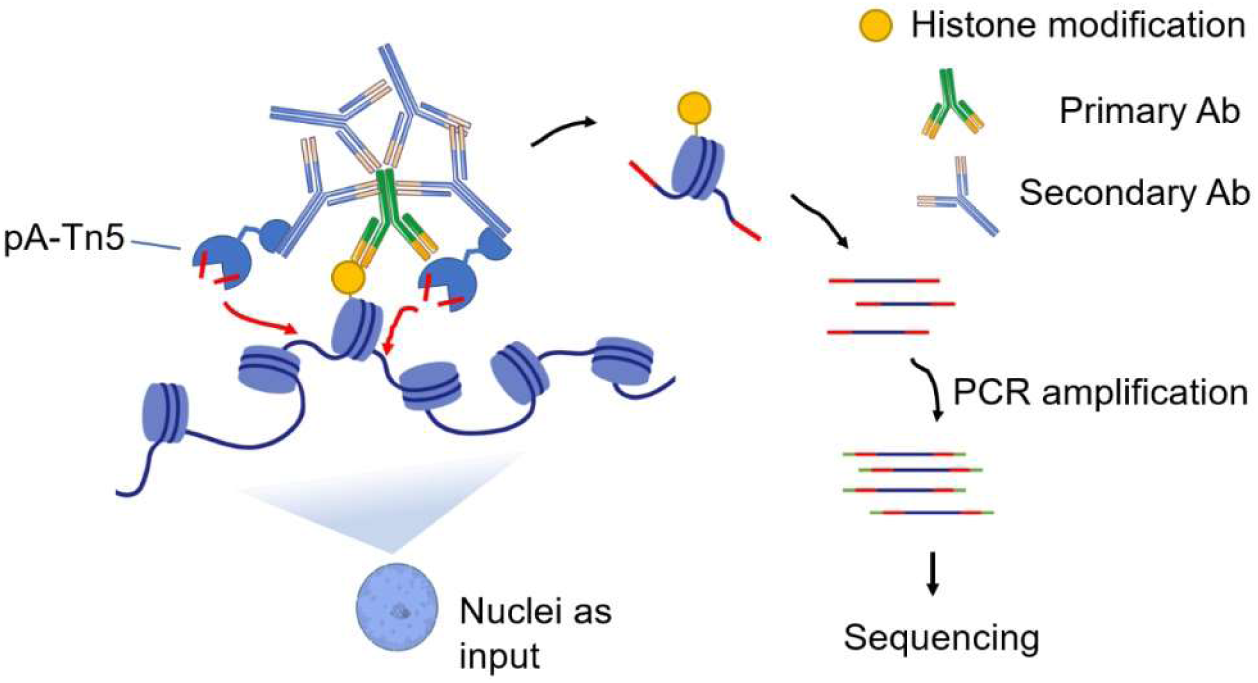
Graphical summary showing the workflow of CUT&Tag using nuclei as input. The nuclei are incubated with a primary antibody that binds to the histone modification of interest. Then, a secondary antibody tethers the pA-Tn5 to the primary antibody. The pA-Tn5 is activated, cleaves and tagments the DNA in proximity. The resulting tagmented DNA is released, and amplified with adapter-containing oligos by PCR, resulting in sequencing-ready libraries.

Compared to ChIP-seq and its variants, CUT&Tag shows better compatibility with extremely low-input materials in animal systems. It has been used with less than 100 mammalian cells, providing high-quality datasets for several histone modifications [1,8]. It was also successfully applied to systems with material limitations, as for example, mammalian embryos at very early stages of development [9].

CUT&Tag represents one of the latest advancements in enzyme-tethering epigenomic profiling techniques. Another popular enzyme-tethering technique is CUT&RUN sequencing (Cleavage Under Targets and Release Using Nuclease)[7], which works in a similar way to CUT&Tag: instead of pA-Tn5, pA-MNase is used to cleavage and release the DNA fragments. Compared to CUT&RUN sequencing, the library preparation for CUT&Tag is more straightforward, because it results from the pA-Tn5 activity and requires only one round of amplification by PCR. In contrast, CUT&RUN requires additional ligation-based library preparation.

Besides its compatibility with small amounts of material, CUT&Tag can also provide a higher theoretical resolution than ChIP-seq and ChIPmentation. In ChIP-seq, the chromatin is shared into 200-300 bp fragments [10]. Therefore, the ends of sequenced fragments may be up to hundreds of base pairs away from the position of the actual target. Differently, in CUT&Tag the transposase is tethered to the target through specific antibodies, thus making the DNA cleavage site in the near proximity of the target. This difference in the distance between the DNA cleavage site and the target should allow CUT&Tag to profile the location of the target most accurately and, therefore, to lead to the highest resolution.

CUT&Tag and CUT&RUN have gained considerable popularity in animal research, and a few reports have already used it to profile the epigenetic landscape in plant tissues [11–20]. However, comprehensive validation of this method is still lacking, raising questions about its ability to replicate ChIP-Seq results, and whether it can provide or not a deeper resolution. In addition, one of the main advantages of CUT&Tag over ChIP-Seq is its requirement for much less starting material. However, this aspect has not yet been investigated in plant tissues. This gap leaves an open question about whether CUT&Tag could expand our portfolio of available techniques, providing a viable alternative for addressing developmental questions in plant research where starting material is often limited. Furthermore, there is no established benchmark or detailed CUT&Tag protocol that reliably works with different antibodies in *Arabidopsis* tissue.

Here, we present an optimized, easy to perform and reliable CUT&Tag protocol adapted to successfully work with *Arabidopsis thaliana* tissues. We tested our protocol with a minimum amount of seedling material (0.01 gram) for three different histone modifications - H3K27me3, H3K4me3 and H3K27Ac – and using histone H3 as a control. By comparing CUT&Tag with ChIP-seq datasets, we show that both CUT&Tag and ChIP-seq capture the same broad lines of the epigenome, while they also capture sets of different peaks. Compared to ChIP-seq, CUT&Tag shows higher sensitivity towards gene bodies, particularly exons. Moreover, we show that CUT&Tag has a higher resolution than ChIP-Seq that reaches the single nucleosomal level. The increased resolution of CUT&Tag allows revealing the detailed patterns of histone modifications occupancy that were hard to decipher in ChIP-seq data, including the enrichment of H3K27me3 at exons, and the epigenetic states of single nucleosomes. These findings prove that CUT&Tag is valid techniques for epigenome profiling in plant tissues that allows to reach deep resolution even with little input material.

## Results

### CUT&Tag reveals similar but also different sets of H3K27me3 peaks compared to ChIP-seq

We adapted and optimized the CUT&Tag protocol, originally developed for mammalian cells, for nuclei extracted from *Arabidopsis* seedlings and tested antibodies from various brands. Our findings indicate that the performance of CUT&Tag can vary significantly depending on the antibody used. Notably, some antibodies effective in ChIP-seq may not be compatible with CUT&Tag (Supplementary Figure 1).

Using the antibodies with best performance, we produced CUT&Tag datasets from *Arabidopsis* wild-type seedlings for the histone modification H3K27me3 using histone H3 as control. To assess the performance of CUT&Tag, we compared our dataset with a publicly available ChIP-Seq dataset (Xiao et al 2017, [21]) and a ChIPmentation dataset (Zhu et al 2023, [6]) for the same histone modification. Starting from a few seedlings, we obtained around 8.7-15.3 million, aligned deduplicated fragments for each H3K27me3 CUT&Tag replicate (Supplementary table 1), which is considered good quality and sufficient for downstream analysis [1]. The H3K27me3 profiles generated by our CUT&Tag analysis show enrichment patterns that typically span one or a few genes. We first noticed that the CUT&Tag H3K27me3 datasets lead to a H3K27me3 occupancy profile similar to the one obtained by ChIP-Seq performed by Xiao et al. (Xiao et al 2017; Fig 2a). Using the same method for peak calling with MACS2, the CUT&Tag H3K27me3 datasets revealed 18’517 peaks that were reproducible in at least two replicates, while 16’858 and 7’842 peaks were detected by Xiao et al. and Zhu et al., respectively. The fewer peaks detected in Zhu et al. suggest that ChIPmentation may be less sensitive than canonical ChIP-Seq, making it more useful for detecting predominantly strong and stable target-DNA interactions. CUT&Tag was able to capture nearly 80% of the ChIPmentation-obtained peaks (Fig 2b), thus suggesting that CUT&Tag is suited to capture robust target-DNA interactions.

**Figure 2.**
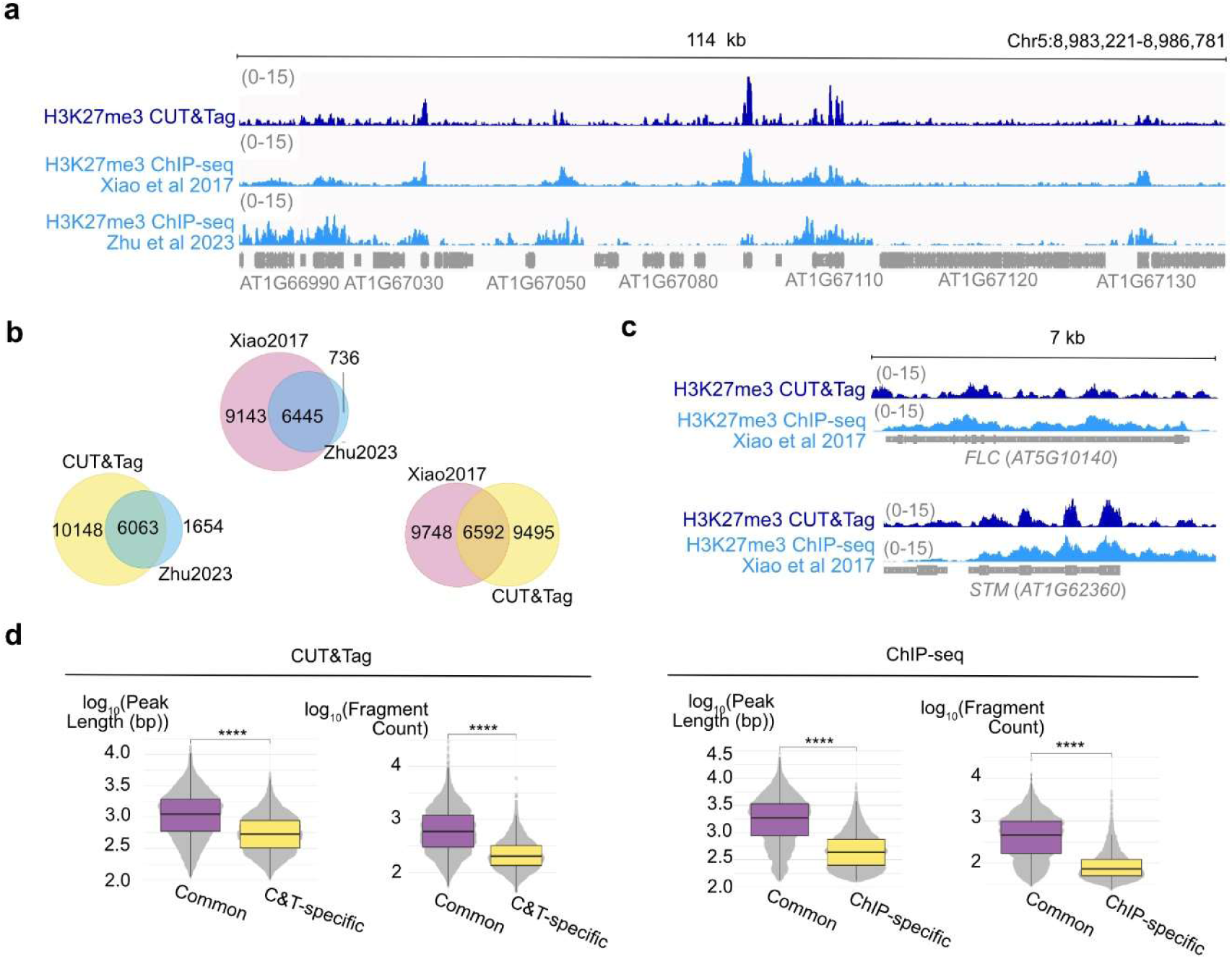
Comparison between ChIP-seq and CUT&Tag. (a) An overview of a 114 kb genomic window, showing the H3K27me3 data from CUT&Tag and two previously published ChIP-seq datasets [6,21]. The profiles show the visual similarity between the H3K27me3 from CUT&Tag and ChIP-seq data from Xiao 2017 et al. (b) Venn diagram depicting the overlap between H3K27me3 peaks found by CUT&Tag (yellow circles) and a ChIP-seq experiment (pink circles) and a ChIPmentation experiment (blue circles). (c) Browser views of H3K27me3 from CUT&Tag and ChIP-seq at two PRC2-target genes showing H3K27me3 enrichment at same locations. (d) Comparisons of the peak length and peak fragment count between peaks common (purple boxplot) in CUT&Tag (left side) and ChIP-seq (right side), and peaks specific (yellow boxplot) to each method. P-value: **** p <= 0.0001, Wilcoxon rank sum test

Next, we focus on the comparison between our CUT&Tag datasets and the ChIP-seq dataset from Xiao et al 2017. While these two datasets share 6’592 common peaks, 9’748 peaks were specific to ChIP-seq (referred to as ChIP-specific peaks), and 9’495 peaks were specific to CUT&Tag (referred to as C&T-specific peaks).

Common peaks were found on genomic location of some known *Polycomb* group proteins target genes, where strong enrichment of H3K27me3 is expected, for example *FLOWERING LOCUS C* (*FLC*) and *SHOOT MERISTEMLESS* (*STM*) (Fig 2c), which are characterized by long peaks (mean length equals 1’463 bp for CUT&Tag peaks and 2’575 bp for ChIP-seq peaks, Fig 2d) with high number of fragment counts (mean fragment count per peak equal 941 for CUT&Tag peaks and 739 fragment counts per peak for ChIP-seq peaks, Fig 2d). The C&T-specific peaks and the ChIP-specific peaks are characterized by short length (mean length equals 672 bp for C&T-specific peaks and 627 bp for ChIP-specific peaks) and fewer read counts than common peaks (Fig 2d), suggesting that these method-specific peaks are associated with less predominant enrichment of H3K27me3.

To assess if the same results would be obtained by a CUT&Tag similar technique, we compared the dataset with a H3K27me3 profiling in leaves obtained by CUT&RUN [17], and found that the CUT&RUN profile of H3K27me3 look visually similar to the CUT&Tag profiles, and has a correlation coefficient between 0.4-0.6 (Supplementary Figure 2). This result suggests that CUT&Tag and CUT&RUN generate similar profiles.

### CUT&Tag resolves the enrichment of histone modifications at exons

To better understand the origin of CUT&Tag and ChIP-seq specific peaks, we plotted their genomic locations, along with the common peaks (Fig 3a). We found that, compared to the common peaks, C&T-specific peaks are more likely located on gene bodies, particularly exons (two examples are shown in Supplementary figure 3), while the ChIP-specific peaks are particularly enriched in the intergenic regions (two examples shown in Supplementary Figure 3).

**Figure 3.**
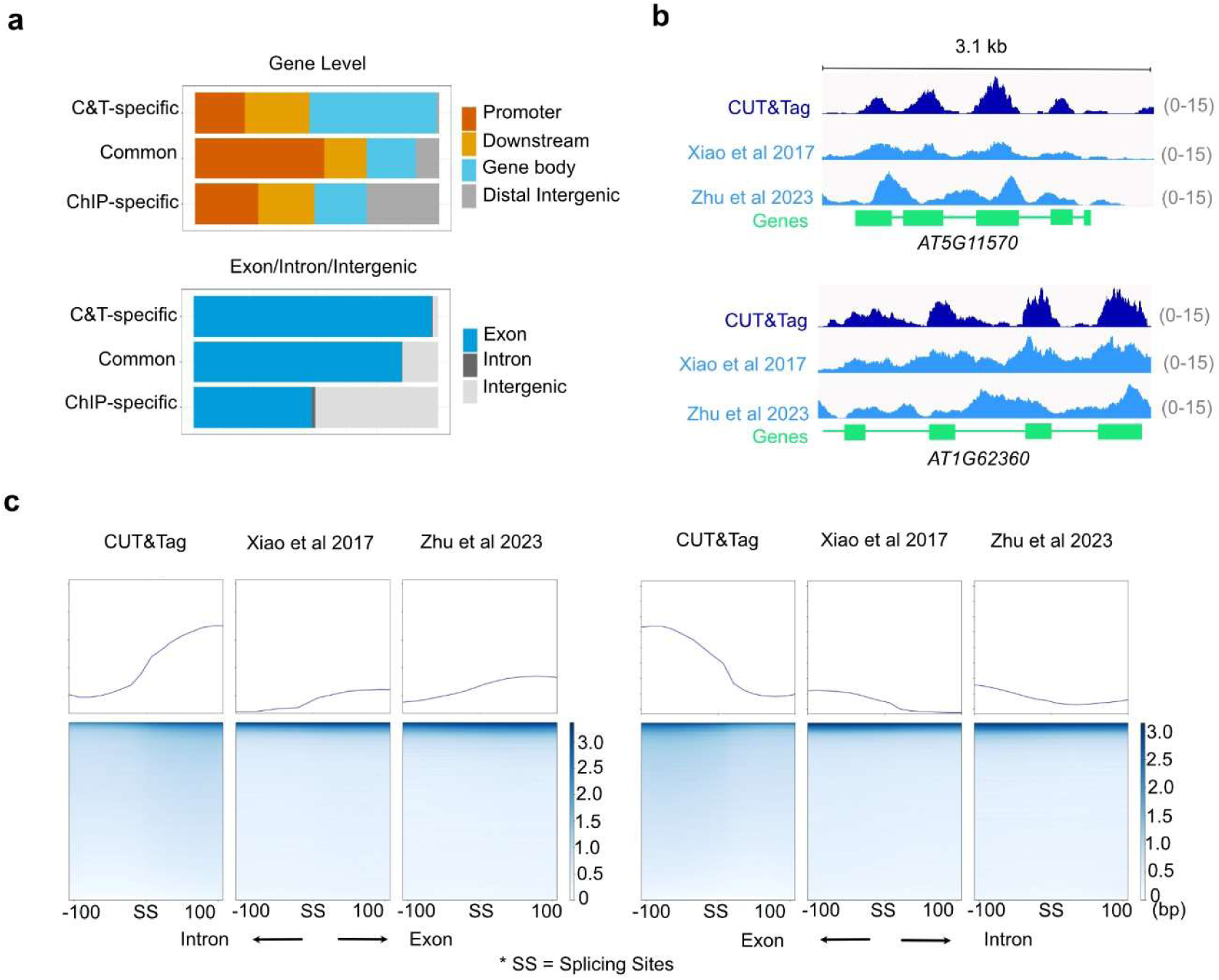
Enrichment of H3K27me3 on exons profiled by CUT&Tag. (a) Comparisons of genomic feature distribution of peaks common in ChIP-seq and CUT&Tag, and peaks specific to each method. (b) Browser view of two example genes showing that CUT&Tag reveals strong enrichment of H3K27me3 on exons compared to introns, in respect to ChIP-Seq and ChIPmentation. (c) Heatplots showing the distribution of H3K27me3 (by CUT&Tag, ChIP-seq and ChIPmentation) around splicing sites.

In line with the observation that CUT&Tag reveals more peaks on exons, we found that the CUT&Tag H3K27me3 signal shows a strong degree of enrichment on exons compared to the introns (Fig 3b). In the ChIP-seq and ChIPmentation data, the exon enrichment was also observed, however not as strong as it was in the CUT&Tag data (Fig 3b). These observations were confirmed with heat plots (Fig 3c).

In summary, we showed that both CUT&Tag and ChIP-seq capture genomic locations stably enriched in H3K27me3 and their specific signatures, thus demonstrating the reliability of CUT&Tag in performing well with *Arabidopsis* tissues. Both techniques capture sets of specific peaks that are short and weak, suggesting that they are both likely affected by minor technical bias. CUT&Tag better reveals the enrichment of histone modifications on exons due to its increased resolution, which leads to the discovery of more exon peaks.

### CUT&Tag profiles of three histone modifications in *Arabidopsis* seedlings

Next, we employed CUT&Tag for profiling two additional histone modifications: H3K4me3 and H3K27Ac, which are two marks that correlates with active transcription. Fig4a provides an overview of the enrichment of these three histone modifications compared to the H3 control, with the peaks indicated. Replicates for each histone modification correlate well and are clustered together (Supplementary Figure 4).

When comparing peak features such as length and location around genes (Fig. 4b, 4c), we noticed that H3K27me3 signals are enriched on gene bodies, while H3K4me3 and H3K27Ac occupancy are particularly enriched around the transcriptional start site (both K4me3 and K27Ac) and gene bodies (mostly K27Ac). Notably, these observations are largely consistent with previous observations obtained by ChIP-seq [23,24].

**Figure 4.**
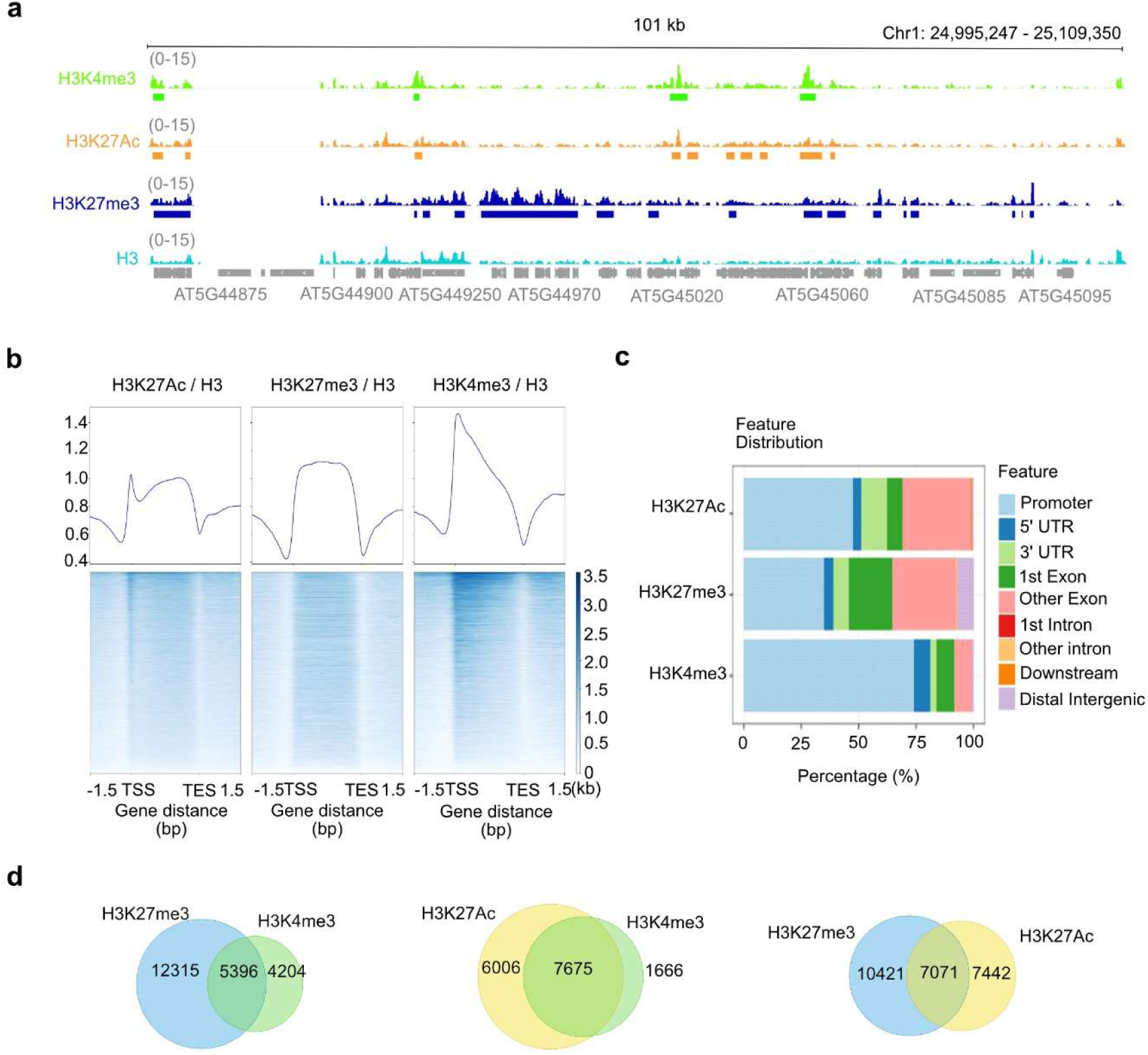
The genome-wide distribution of three histone modifications profiled by CUT&Tag. (a) A browser view of 101-kb genomic windows showing the profiles of three histone modifications and the H3 control profiled by CUT&Tag. The profiles show different features of peaks of the three histone modifications: H3K4me3 peaks are mostly located around transcription start sites, H3K27Ac peaks are located at promoters and gene bodies, and H3K27me3 peaks usually span one or a few genes. (b) Heatplot showing the distribution of H3K27me3, H3K4me3 and H3K27Ac around the gene bodies. Abbreviations: TSS, transcriptional start sites; TES, transcription end sites. (c) Genomic feature distribution of H3K27me3, H3K4me3 and H3K27Ac peaks. (d) Venn plots showing the pairwise overlap of CUT&Tag peaks of H3K27me3 (blue circle), H3K4me3 (green circle) and H3K27Ac (yellow circle).

We then compared the H3K27me3, H3K4me3 and H3K27Ac datasets, and looked for common and different peaks (Fig 4d). As expected, H3K27Ac and H3K4me3 peaks shared a high degree of overlap, as they both have the same activating effect on transcription. Interestingly, we found that around half of the H3K27Ac and H3K4me3 peaks are also marked by H3K27me3. This degree of overlap is higher than the previously reported 15% overlap between H3K27me3 and H3K4me3 [25].

### CUT&Tag reveals the relations between histone marks and gene expression

To understand if the epigenetic status revealed by CUT&Tag correlates with the transcriptional status of the peak-bearing genes, we compared our CUT&Tag datasets with publicly available transcriptomic data (RNA-Seq) from *Arabidopsis* seedlings. We observed that the presence of a certain histone modification correlates well with the expression levels of the targets detected in seedlings. For instance, floral and meristem identity genes such as *APETALA1* (*AP1*) and *STM* showed strong enrichment of H3K27me3 and depletion of H3K4me3 and H3K27Ac. Accordingly, both *STM* and *AP1* are repressed in seedlings by the activity of *Polycomb* group proteins [6,21,22](Supplementary Figure 5a). Actively transcribed genes, such as *RIBOSOMAL PROTEIN L7B* (*RPL7B*, *AT2G01250*) and *UBIQUITIN CARRIER PROTEIN 1* (*UBC1*, *AT1G14400*), exhibited enrichment of H3K4me3 and H3K27Ac (Supplementary Figure 5b).

Next, we plotted the expression levels of genes marked and not marked by each histone modification (Fig 5a), where a gene is classified as marked if at least one peak is detected in its promoter, 5’UTR, gene body, or 3’UTR regions. H3K4me3-marked and H3K27Ac-marked genes showed significantly higher expression than not-marked genes, which is consistent with the known association of these histone modifications with active transcription [26]. Interestingly, we found that H3K27me3-marked genes had a slightly higher average expression level than unmarked genes, which is not the expected correlation as H3K27me3 is a mark associated with gene repression [27]. This could be explained by the fact that we used whole seedlings as starting material, which consist of various cell and tissue types. As a result, the dataset represents a mixture of occupancy from different cell types, where certain genes may be marked differently. Therefore, many of the H3K27me3 genes are likely only marked by H3K27me3 in a small population of cells but actively expressed in other cells. To further explore the relation between H3K27me3 and gene expression, we quantified the H3K27me3 enrichment level (H3K27me3 / H3) and found that genes that are highly enriched in H3K27me3 tend to have low expression levels (Fig 5b). In agreement, we found that genes normally expressed in discrete domains of the seedlings, have both H3K27me3 and H3K4me3 marks. For instance, *TEOSINTE BRANCHED1/CYCLOIDEA/PCF 15* (*TCP15*, AT1G69690) was enriched in both H3K4me3/H3K27Ac and H3K27me3 modifications (Supplementary Figure 5c). TCP15 is normally expressed in young, proliferating leaves but silenced in older ones [28]. Therefore, the detected mixed chromatin state may be the result of the sum of individual chromatin states, repressed and active, that depends on the cell type. This scenario became even more evident when comparing all the three histone modifications against each other (Fig 5c). Indeed, we observed that around half of the H3K4me3/H3K27ac marked genes are also marked by H3K27me3.

**Figure 5.**
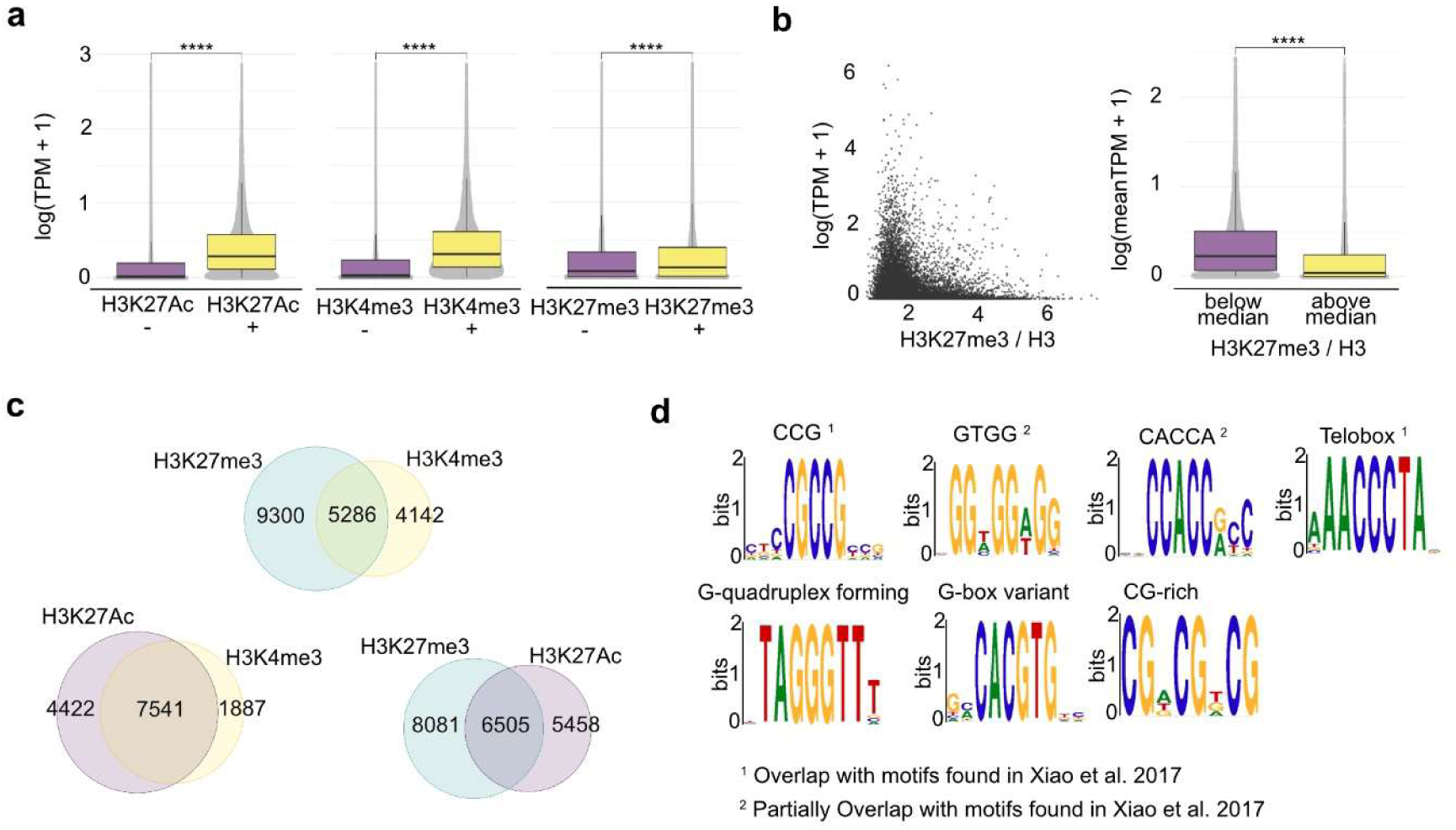
Relation between histone marks and gene expression level, overlap between different histone marks, and motif enrichment analysis using the CUT&Tag datasets. (a) Bar plots showing the comparison of the expression level between genes not marked (purple boxplot) and marked (yellow boxplot) by each histone modification. (b) Dot plot and bar plot showing the anti-correlation between the H3K27me3 level and expression among all H3K27me3-marked genes. (c) Venn diagram showing the pairwise overlap of genes marked by different histone marks H3K27me3 (blue circles), H3K4me3 (yellow circles) and H3K27Ac (purple circles). (d) Enriched motifs found from all replicates of H3K27me3 peaks; the motifs that are overlapped or partially overlapped with a previous report (Xiao et al. 2017) are labelled by 1 and 2 P-value: **** p <= 0.0001, Wilcoxon rank sum test

In summary, we utilized CUT&Tag to map histone modification occupancy and coupled this with transcriptome analysis, providing biological insights into the effects of specific epigenetic states on gene transcription.

### Motif enrichment analysis using CUT&Tag data reveals motifs potentially involved in H3K27me3 deposition

Deposition of histone marks is made by histone modifying-enzyme, which are recruited at genomic loci by the interaction with other proteins, or also by consensus sequences. In *Arabidopsis*, the recruitment of the *Polycomb* Repressive Complex 2 (PRC2) is likely through interaction of the complex with transcription factors that binds to cis elements in the genome. Therefore, a motif search analysis of H3K27me3-marked genomic fragments might help revealing cis elements that favor PRC2 recruitment [21]. The motif enrichment analysis on H3K27me3 CUT&Tag peaks revealed seven motifs (Fig 5d). Notably, four out of seven motifs overlap or partially overlap with motifs previously showed to be associated with PRC2 recruitment (Xiao et al. 2017), including for instance the Telobox. The other motifs, including a G-quadruplex forming sequencing [29] and a G-box variant [30], have no known connections with H3K27me3, and a CG-rich motif, have not been reported before. To conclude, we explored DNA motif enrichment within the CUT&Tag datasets, which could aid in identifying chromatin regulators with role on histone modifications deposition/removal.

### CUT&Tag has an increased resolution compared to ChIP-seq

Histone modification occupancy obtained by CUT&Tag showed a wavy pattern, with bulges of around 200-300 bp (Fig 6a), that corresponds to the length of the DNA wrapped around a nucleasome. To assess if indeed CUT&Tag can reach nucleosome-size resolution, we compared the CUT&Tag data with a public MNase-seq data of *Arabidopsis* seedlings [31]. We examined the H3K27me3 signal around two genes which are H3K27me3-marked in seedlings, *FLC* and *MEDEA* (*MEA*), and observed indeed that the CUT&Tag-detected H3K27me3 signal and MNase-seq showed overlapping wavy pattern in many regions (Fig 6a). This clear pattern could not be captured by neither ChIP-Seq (Xiao et al 2017, [21]), or ChIPmentation (Zhu et al 2023, [6]) where the peaks are averaged with the neighboring regions and lack details. Notably, we could observe a similar pattern from CUT&RUN data [7] (Supplementary Figure 6). However, CUT&Tag tends to have a lower background in the inter-nucleosome regions, and the individual nucleosomes are more recognizable with CUT&Tag data than with CUT&RUN. Thus, CUT&Tag provides nucleosome-level resolution of histone modification occupancy, at a depth that is unmatched by other profiling techniques. Interestingly, we noticed that, in some cases the two adjacent nucleosomes could have quite different levels of H3K27me3 (Fig 6a, indicated by red arrows). Moreover, we observed clear epigenetic states for the +1 nucleosome (Figure 6b), which is involved in transcription elongation, pre-mRNA processing and gene expression [32]. In some cases, the +1 nucleosomes were marked by H3K27me3 (Figure 6b left), while in some other cases, they are marked by active marks H3K4me3 and H3K27Ac (Figure 6b right). These interesting observations show that CUT&Tag can be used to investigate the epigenetic states of individual nucleosomes due to its increased resolution.

**Figure 6.**
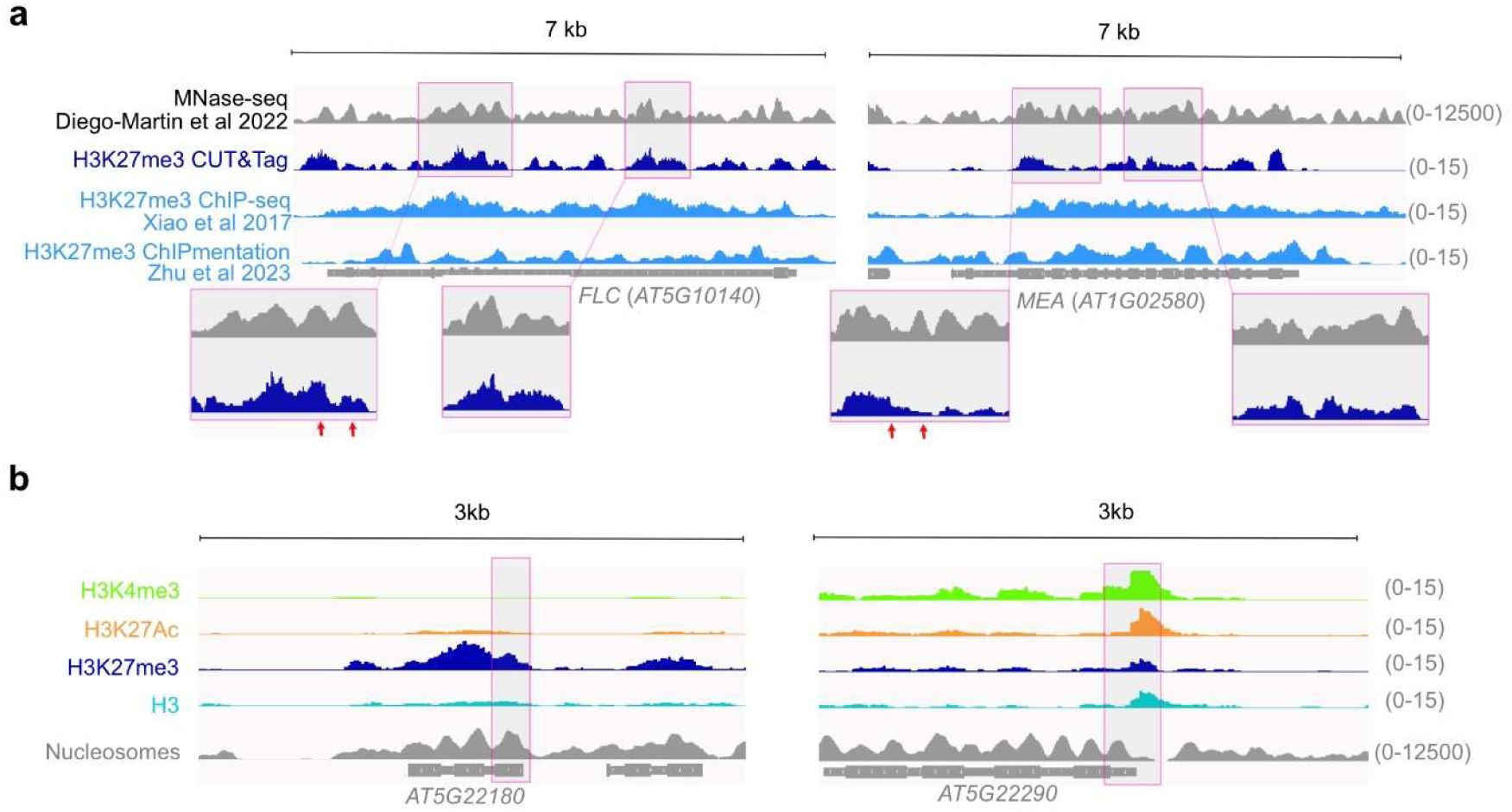
Functional analysis using CUT&Tag datasets. (a) Genome browser view showing the comparison between the nucleosome occupancy (MNase-seq, [31]) and H3K27me3 profiles from CUT&Tag and two ChIP-seq datasets; Regions in the CUT&Tag dataset show good cooreltion with the nucleosome occupancy are highlighted by a grey box; adjacent nucleosomes with strongly different H3K27me3 levels are pointed by red arrows. (b) Genome browser view of two genes showing the epigenetic states of the +1 nucleosome profiled by CUT&Tag.

## Discussion

Here, we present an optimized and validated protocol for CUT&Tag targeting three different histone modifications in the tissues of the model plant *Arabidopsis thaliana*. This protocol primarily follows the steps outlined in previously published CUT&Tag protocols for mammalian cells [1,8]. Here. we have only optimized it to work efficiently with plant nuclei, and have also tested antibodies from various brands. Our results indicate that some antibodies which perform well in ChIP-seq may not work as well in CUT&Tag (Supplementary Figure 1). Therefore, we recommend using antibodies specifically tested for CUT&Tag when replicating these experiments.

By comparing our CUT&Tag-generated datasets, with two publicly available ChIP-seq datasets, we showed that CUT&Tag is a valid method for epigenome profiling, and explored the differences and the advantages of CUT&Tag versus ChIP-seq. Indeed, CUT&Tag captures the precise genome-wide distribution of three histone modifications: H3K27me3, H3K4me3 and H3K27Ac from a small amount of material, and also reaches nucleosomal resolution, which neither ChIP-seq nor ChIPmentation can achieve.

ChIP-seq and CUT&Tag are orthogonal methods for epigenome profiling, and there have been some reports stating that they reveal some different sets of peaks [1,14,16,33]. In our work, we also found that CUT&Tag and ChIP-seq both captured common but also sets of peaks specific to themselves. The ChIP-seq specific peaks are enriched in intergenic regions with low reads density. Considering that ChIP-seq is known to produce low-density genome-wide noise [7], many of the intergenic ChIP-specific peaks are likely noise from low-signal regions.

We also showed that the CUT&Tag-specific peaks are enriched on exons, suggesting CUT&Tag has an increased sensitivity for the latter. We think this is a result of the increased resolution of CUT&Tag. The average lengths of exons and introns of *Arabidopsis* are around 250 and 168 bp, respectively [34,35]. The lower resolution of ChIP-seq hardly resolves the precise distribution of histone modifications on introns and exons, and therefore specific enrichments in exons might been hidden withing much longer peaks. Although it has been known that nucleosomes and some histone modifications are more enriched in exons than introns, which may instruct splicing [36,37], the enrichment of H3K27me3 on *Arabidopsis* gene exons has not been previously reported yet. Given H3K27me3’s significant role in gene repression and development, investigating its impact on splicing and its interaction with known functions of H3K27me3 would be valuable. Also, the degree of exon enrichment of H3K27me3 seems to be stronger in *Arabidopsis* compared to that in animal cells [38], possibly suggesting an interesting crosstalk between splicing and chromatin structure in *Arabidopsis*, which could be a interesting avenue of research.

We observed weak enrichment of the histone modification H3K27me3 on the gene bodies of many genes, including housekeeping genes, which are actively expressed in seedlings and should not be marked by H3K27me3. We attribute this observation to the increased sensitivity of CUT&Tag, which allows for the detection of even weak signatures of H3K27me3 enrichment within exons. Moreover, given that our starting material - the *Arabidopsis* seedling - consists of a mixture of different cell types and that epigenetic states are cell type-specific, this weak enrichment might come from H3K27me3 occupancy in a small subset of cells. Therefore, although CUT&Tag performed on bulk tissues like seedlings does not directly reveal the epigenetic states of individual cell types, we propose that combining this technique with gene expression data could provide insights into the epigenetic landscape across different cell types. More tailored studies are needed to correlate gene expression patterns with observed histone mark enrichment, thereby inferring the epigenetic states within distinct cell populations.

We showed that an advantage of CUT&Tag versus ChIP-seq is its compatibility with small amounts of starting material. In this work, we used just a few one- or two-week-old seedlings for each reaction, which was about 0.01 grams instead of several grams required by ChIP-seq. By this, CUT&Tag can enable research with limits in material availability. For example, the research of epigenetic variation between individual plants or the research of epigenetic states of a certain tissue types during flower development. Furthermore, we obtained informative and robust data and sequencing depth from seedlings, and we thus suggest that it would be possible to have good quality datasets with even less amount of material.

We demonstrated that CUT&Tag achieves single-nucleosome resolution, thus providing an accurate depiction of the epigenetic status of a gene. This is a significant improvement in respect to ChIP-seq, where nucleosome patterns are often ambiguous and histone modifications are generally detected spread across long genomic regions. Our CUT&Tag data revealed that even adjacent nucleosomes within H3K27me3-enriched regions can exhibit markedly different levels of H3K27me3. This variation in epigenetic mark levels at the single-nucleosome level presents an intriguing area for further investigations. Additionally, we observed distinct epigenetic states for the +1 nucleosome., with the +1 nucleosome peaks in CUT&Tag data shifted towards the nucleosome depletion region (Fig. 6b, right). This could be due to nucleosome depletion regions attracting tagmentation by the transposase tethered to the +1 nucleosomes, or a nucleosome covered by transcription-related proteins that MNase-seq can not resolve. Thus, researchers studying the epigenetic states of +1 nucleosomes could greatly benefit from this technique.

## Conclusions

We presented a CUT&Tag protocol for the model plant Arabidopsis. CUT&Tag captures the genome-wide distribution of histone modifications while using much less material than ChIP-seq. Additionally, CUT&Tag can reach a nucleosome-level resolution.

## Authors’ contributions

YF performed the experiments and analysed the data; MWS contributed with data analysis and provided assistance and feedback; SS provide supervision, analysed the data and acquired fundings; YF and SS wrote the manuscript, with MWS commenting on it.

## Availability of data and materials

Raw sequencing data are available at NCBI under the accession number XXXXXxxxxxxx.

## Acknowledgement

We thank Prof. Ueli Grossniklaus (University of Zurich) for sharing equipment; Peter Kopf (University of Zurich) for general lab support; Elizabeth Kracik-Dyer (University of Zurich) and Dr. Célia Baroux (University of Zurich) for helping with the nuclei extraction; the Functional Genomics Center Zurich for providing the high-throughput sequencing service; Dr. Moccia Maria Domenica (Functional Genomics Center Zurich), Dr. Susanne Kreutzer (Functional Genomics Center Zurich) for helpful suggestions about sequencing; Prof. Steve Henikoff (Fred Hutchinson Cancer Research Center) for generating and depositing the 3XFlag-pA-Tn5-Fl plasmid; and the CHERI T meeting (University of Zurich) for useful discussion.

## Methods

### Plant material and growth conditions

*Arabidopsis thaliana* ecotype Col-0 (Columbia) NASC #N70000 seeds were sown on half-strength MS media (1/2 MS salt base [Carolina Biologicals, USA], 1% sucrose, 0.05% MES, 0.8% Phytoagar [Duchefa], pH>5.7 with KOH) and subjected to stratification at 4°C for 2-3 days. Following this, the plates were moved to a plant incubator (CU36L6, CLF Plant Climatics) set to long-day conditions (8h dark at 18°C, 16h light at 22°C, 70% relative humidity). When harvesting seedlings, the plants were grown on half-strength MS media for between one or two weeks, allowing for the development of 4-8 true leaves.

### Antibodies used in this study

Rabbit anti-H3K27me3: Cell signaling technology #9733

Rabbit anti H3K4me3: Diagenode c15410003

Rabbit anti H3K27Ac: Abcam #4729

Mouse anti-H3: Active Motif 39064

Guinea Pig anti-Rabbit IgG: antibodies-online #ABIN101961

Rabbit anti-Mouse IgG: Abcam 46540

### Production of pA-Tn5

The procedure for purifying pA-Tn5 is based on previously reported protocols [1,39] and the workflow is summerized in Supplementary Figure 7a.

HEGX buffer and dialysis buffer were prepared before starting the protein production. HEGX buffer is prepared using the following recipe: 20 mM HEPES pH7.2 [Sigma-Aldrich #H3375], 1 M NaCl [Roth #3957.1], 1 mM EDTA pH8 [HUBERLAB # A2937.1000], 10% Glycerol [Roth #3783.1], 0.2% Triton X-100 [Sigma #T8787], Protease Inhibitor (1X, add freshly before use) [Roche #64178100]. Dialysis buffer is prepared using the following recipe: 100 mM HEPES pH7.2 [Sigma-Aldrich #H3375], 200 mM NaCl [Roth #3957.1], 0.2 mM EDTA [HUBERLAB # A2937.1000], 20% Glycerol [Roth #3783.1], 0.2% Triton X-100 [Sigma #T8787], 2 mM DTT [Merck # 1.24511.0005].

*E.coli* containing the plasmid the 3XFlag-pA-Tn5-Fl [Addgene Plasmid #124601, C3013 *E. coli* strain] was obtained from Addgene, and propagated on plate by streaking the bacteria from the agar stab onto an LB agar plate supplemented 100 μg/mL Ampicillin [Applichem #A0839] and let to grow overnight at 37°C. A single colony was then inoculated in 7.5 mL of liquid LB medium supplemented with 100 μg/mL Ampicillin and let to grow at 37°C, shaking at 180rpm for about 4 hours. This 7.5 mL of culture was then added to 500 mL of liquid LB medium supplemented with 100 μg/mL Ampicillin and left to grow at 37°C, shaking 180rpm until an OD600 value between 0.5 and 0.7 was reached. Then, the liquid culture was cooled down on ice for one hour, and 125 μL of 1M IPTG (final concentration 0.25 mM) were added to induce the expression of the protein, overnight at 18°C shaking at 180rpm. The following morning, the colture was centrifuged at 9500 x g at 4°C for 1h to pellet the bacteria using a Sorvall LYNX 6000 Superspeed Centrifuge (Thermo Scientific #75006590), the supernatant removed, and the bacteria pellet frozen -80°C for at least two hours. To purify the protein, the bacteria pellet was thawed on ice, resuspended in 5 volumes of HEGX buffer supplemented with 0.5 mg/mL Lysozyme [Roche #10837059001], and incubated on ice for 30 minutes. After the incubation, the bacteria suspension was sonicated with a Bioruptor bath sonicator (Diagenode), with setting M, 15s on, 15s off, 10 cycles. The lysate was then centrifuged at 31,000 x g, 4°C, for 30 minutes and the supernatant mixed with 5 mL Chitin Resin (NEB #S6651), previously washed first with 30 mL of ddH2O and then 20 mL of HEGX buffer. The samples were incubated at 4°C for 48 hours rotating. After incubation, the sample was transferred into a 10 mL gravity column (Thermo Scientific #89898). Once all the resin was packed in the gravity column, the resin was washed with 20 mL HEGX buffer supplemented with protease inhibitor (1X, Roche #64178100), and then the protein eluted with elution buffer (HEGX buffer with 100 mM DTT [Merck # 1.24511.0005]). The protein prep was then dialyzed with SnakeSkin Dialysis Tubing (3500 MWCO, Thermo Scientific #68035) using 800 mL of dialysis buffer at 4°C. The quality of the purified pA-Tn5 protein was confirmed by SDS-PAGE followed by coomassie blue staining and its concentration was estimated by comparing to BSA standards on the same gel. Then, the protein was concentrated by centrifugation with a Pierce Protein Concentrator (10K MWCO, Thermo Scientific #88516) to a concentration close to 800 ng/μL. An equal volume of filter-sterilized 80% Glycerol (Roth #3783.1) was added to the protein prep, bringing the protein concentration to about to 400 μg/mL (about 5.5 μM). The protein was aliquoted, and aliquots stored at -20°C.

DNA oligos used to create the adaptors including ME-A (Sequence: TCGTCGGCAGCGTCAGATGTGTATAAGAGACAG), ME-B (Sequence: GTCTCGTGGGCTCGGAGATGTGTATAAGAGACAG), and MErev (Sequence: /5Phos/CTGTCTCTTATACACATCT) were synthetized with standard desalt purification and resuspended in annealing buffer (10 mM Tris-HCl pH 8.0, 50 mM NaCl, 1 mM EDTA) to create a stock concentration of 200 mM. To form ME-A and ME-B adaptors, the pair ME-A plus MErev (40 μL each), and ME-B plus MErev (40 μL each) were mixed, heated at 95°C for 5 minutes, and left to cool to room temperature in the heating block turned off. The formed ME-A and ME-B adaptors were then stored at 4°C. To load pA-Tn5 with the adaptors, 4 μL ME-A adaptor, 4 μL ME-B adaptor, and 50 μL pA-Tn5 (at approximately 400 μg/mL) were mixed and incubated at room temperature for 50 minutes and then stored at - 20°C.

To test the transposase activity of the loaded pA-Tn5, 1.25 μL of loaded pA-Tn5 were added to 1 μL of a plasmid at 250 ng/μL (we used a derivative of pAGM65879 [Addgene #153214]), 1 μL NEBuffer 3.1 (NEB #B6003S), 6.75 μL ddH2O, and incubated at 37°C for 15 minutes. To stop the reaction, 1 μL of 1% SDS (Roth #CN30.3) was added, the sample incubated at 55°C for 10 minutes, and then 0.32 μL Proteinase K (Thermo Scientific #EO0492) were added, and the sample incubated further at 55°C for 10 minutes. Complete digestion of the plasmid, as expected if pA-Tn5 has a good transposase activity, was evaluated by loading the sample on 1% Agarose gel (Supplementary Figure 7b, 7c).

The loaded and unloaded pA-Tn5 are stored at -20°C. Please note that the purified pA-Tn5 stock contains a trace amount of *E. coli* DNA, and the loaded pA-Tn5 can slowly tagment the *E. coli* DNA even if stored at -20°C, leading to a increased tagmented *E. coli* DNA level and a decrease of avtivity (Supplementary Figure 8, [1]). Therefore, the tagmented pA-Tn5 should not be stored for more than six months.

### Reagent Setup for Nuclei Extraction

#### LB01 Buffer for nuclei extraction ( 40mL)

0.6 mL 1M Tris-HCl, pH 8 [Roth # AE15.3] (final: 15 mM)

0.16 mL 0.5M EDTA [HUBERLAB #A2937.1000] (final: 2 mM)

20 μL 1M Spermidine [Spermidine trihydrochloride: Sigma-Aldrich #S2876] (final: 0.5 mM)

1.6 mL 2M KCl [Roth #6781.1] (final: 80 mM)

0.16 mL 5M NaCl [Roth #3957.1] (final: 20 mM)

0.8 mL 5% Triton [Sigma #T8787] (final: 0.1%)

36.66 mL ddH2O

Store at 4°C for short term uses. Make 10 mL aliquots and freeze at -20 °C for long term storage. 1 mL LB01 buffer is used for nuclei extraction from each seedling sample contained in one 1.5-mL tube.

Add 1/25 volume of Protease Inhibitor [cOmplete, EDTA-free Protease Inhibitor Cocktail, Roche #6417810] (25X) before use.

#### Honda Buffer for nuclei extraction (40 mL)

1.6 mL 0.5M HEPES, pH 7.5 [Sigma-Aldrich #H3375] (final: 20 mM)

17.6 mL 1M Sucrose [PanReac AppliChem #A2211] (final: 0.44 M)

6.2 mL 8% Ficoll PM400 [Merck #F437] (final: 1.25%)

8 mL 12.5% Dextran T-40 [Roth #7626.4] (final: 2.5%)

0.4 mL 1M MgCl2 [MgCl2*6H2O Sigma-Aldrich # M2670] (10 mM)

4 mL 5% TritonX-100 [Sigma #T8787] (final 0.5%)

0.2 mL 1M DTT [Merck # 1.24511.0005] (final: 5 mM)

0.4 mL ddH2O

Store at 4°C for short-term uses. Make 5 mL aliquots and freeze at -20 °C for long term storage. 1 mL LB01 buffer is used for nuclei extraction from each seedling sample contained in one 1.5-mL tube.

1/25 volume of Protease Inhibitor [cOmplete, EDTA-free Protease Inhibitor Cocktail, Roche #6417810](25X) before use

### Nuclei extraction from seedlings

The aerial parts of two to four seedlings between one and two weeks old were collected to extract the nuclei for each CUT&Tag reaction. The seedlings were collected into a 1.5 mL DNA LoBind tube (Eppendorf #L203809K). The tubes containing the sample were snap frozen in liquid nitrogen, and then ground with a Silamat S6 tissue grinder (or similar) for 5 seconds while the tissue was still frozen. The freeze-grind cycle was repeated for a total of three times for each tube. After grinding, 800 μL of Honda buffer was added to each tube, and the sample was then incubated on ice for 20 minutes. The sample was then filtered through a 30 μm CellStrainer (CellTrics) into a clean 2 mL centrifuge tube. An additional 600 μL Honda buffer was used to wash the 1.5 mL tube and the Cell Strainer and merged to the initial 800 μL in the appropriate 2 mL sample tube. A final volume of approximately 1.4 mL was obtained at this step. The nuclei samples were then centrifuged at 4000 x g for 15 minutes at 4°C, and then the supernatant was removed. To wash the nuclei pellet, 1 mL LB01 buffer was added. The tubes were then centrifuged again at 1000 x g at 4°C for 15 minutes, and the supernatant was removed. The nuclei pellet was then resuspended in 200 μL of Wash Buffer. This step uses an EDTA-containing buffer (LB01) to remove the residual Mg2+ from the Honda buffer, and then the nuclei are resuspended in an EDTA-free buffer (Wash buffer). The nuclei quality was checked using a microscope before proceeding with the CUT&Tag.

### Nuclei quality check

For quality control, 3 μL of the nuclei suspension was mixed with 3 μL of VECTASHIELD Antifade mounting medium supplemented with DAPI (Vector Laboratories, H-1200). The sample was loaded on a Neubauer hemocytometer (VWR #631-0926), and the nuclei integrity and number were assessed with a Leica DM6000 fluorescence microscope. The nuclei of Arabidopsis have diameters between 1 to 10 μm, generally with a smooth contour. A nucleus can be recognized by its chromocenters and sometimes the nucleolus. A good nucleus may have 1 – 10 chromocenters.

### Reagent Setup for CUT&Tag

#### Binding buffer for activating Con-A beads (10 mL)

400 μL 0.5M HEPES [Sigma-Aldrich #H3375], pH 7.9

50 μL 2M KCl [Roth #6781.1]

10 μL 1M CaCl2 [Sigma-Aldrich #21074]

10 μL 1M MnCl2 [Merck # 1.05927.0100]

9.5 mL ddH2O

Store at 4°C for up to 6 month

#### Wash Buffer (50mL)

2 mL 0.5M HEPES [Sigma-Aldrich #H3375], pH 7.5 (final: 20 mM)

1.5 mL 5M NaCl [Roth #3957.1] (final: 150 mM)

25 μL 1M Spermindine [Spermidine trihydrochloride: Sigma-Aldrich #S2876] (final: 0.5 mM)

2 mL 25X Protease Inhibitor Cocktail [cOmplete, EDTA-free Protease Inhibitor Cocktail, Roche #6417810]) (final: 1X)

Add ddH2O to a final volume of 50 mL, filter sterilize

#### Tween-Wash Buffer (20 mL)

550 μL needed for each reaction; prepared by adding 400 μL 5% Tween 20 [Sigma #P9416] (final: 0.1%) to 20 mL Wash buffer

#### Antibody buffer (1 mL)

50 μL needed for each reaction; prepared by adding 4 μL 0.5M EDTA [HUBERLAB #A2937.1000] (final 2 mM) and 10 μL 10% BSA [Sigma-Aldrich #A4503] (final 0.1%).

1 mL Tween-Wash buffer

#### 300-Wash Buffer (50 mL)

2 mL 0.5M HEPES [Sigma-Aldrich #H3375], pH 7.5

3 mL 5M NaCl [Roth #3957.1] final 300 mM

25 μL 1M Spermindine [Spermidine trihydrochloride: Sigma-Aldrich #S2876]

2 mL 25X PI [cOmplete, EDTA-free Protease Inhibitor Cocktail, Roche #6417810]) (final: 1X)

ddH2O to a final volume of 50 mL, filter sterilize

*300 stands for the concentration of NaCl in the buffer

#### Tween-300-Wash Buffer (20 mL)

760 μL needed for each reaction; prepared by adding 40 μL 5% Tween 20 [Sigma #P9416] (final: 0.01%) to 20 mL of 300-Wash buffer

#### Tagmentation buffer (4 mL)

300 μL needed for each reaction; 4 mL Tween-300-wash buffer + 40 μL 1M MgCl2 [MgCl2*6H2O Sigma-Aldrich # M2670] (final: 10 mM).

### CUT&Tag

(Day 1)

#### Preparing Concavalin-A (Con-A) Beads

1. Resuspend Concavalin-A beads [BioMag Plus Concanavalin A, Polyscience Inc, #86057-10] by shaking the bottle. Transfer enough volume to a 2 mL tube containing 1.6 mL of Binding Buffer and incubate for 2 minutes. 15 μL of Con-A beads was used for each sample.
2. Put the tube on a magnet stand [Invitrogen #CS15000] and wait until the supernatant becomes clear (takes 1-2 minutes). Then, remove the supernatant using a 1000 uL pipette.
3. Resuspend the beads using another 1.5 mL Binding Buffer by pipetting up and down. Then, put the tube on the magnet stand, wait until clear, and remove the supernatant.
4. Pipette to resuspend the Con-A beads with Binding buffer, the volume of the binding buffer to use here is the same as the volume of Con-A beads used in step 1. The Con-A beads are now activated.

#### Binding of Nuclei to Con-A Beads, and Binding of Primary Antibody

5. Dispense 15 μL of the activated Con-A beads to each sample of nuclei (which are in the Wash buffer). Rotate the mixture of the nuclei and beads at room temperature for 10 minutes.
6. Meanwhile, dilute the primary antibody of choice in the Antibody buffer. For each sample, prepare 50 μL of Antibody buffer, and add antibody at a dilution ratio of 1:50, or follow the dilution ratio suggested by the antibody provider or previous reports.
7. After the 10 minutes of incubation (step 5) are passed, put the tubes on the magnet stand, wait until the supernatant is clear, then remove the supernatant with a P200 pipette. Immediately resuspend each sample in the 50 μL Antibody buffer prepared in step 6.
8. Put the samples to rotate at 4°C and incubate overnight.

(Day 2)

#### Binding of Secondary Antibody

9. Dilute the secondary antibody of choice in Tween-Wash Buffer (55 μL for each sample).
10. After the overnight incubation of the sample, quickly spin the tube with a microcentrifuge to collect all the liquid on the bottom of the tube, then place the tubes on the magnet stand.
11. Using a P200 pipette set at 55 μL, remove the supernatant, then immediately add 55 μL of the dilution of the secondary antibodies.
12. Resuspend the beads by gently vortex the tube and incubate rotating at room temperature for 45 minutes.

#### Binding of the pA-Tn5

13. Dilute the pA-Tn5 (Loaded with adaptors, 4.74 uM) at 1:250 in Tween-300-Wash buffer. Note: home-made pA-Tn5 contains *E. coli* DNA, which serves as spike-in DNA that can be used for normalization [1].
14. After the 45 minutes incubation (step 3.4), briefly spin the samples, put them on a magnet stand and use a P200 pipette set at 60 μL to completely remove the supernatant. Then, while the tubes stay on the magnet stand, add 500 μL Tween-wash buffer to each tube.
15. Remove the majority of the supernatant first with the P1000 pipette, then the remaining with P200 pipette. If there is liquid attached to the wall of the tube, briefly spin the tube to collect the liquid at the bottom of the tube and then completely remove it with the P200 pipette.
16. Add 60 μL pA-Tn5 dilution and resuspend by gently vortex the tube.
17. Rotate at room temperature for 1 hour.

#### Tagmentation

18. After the incubation (step 17), briefly spin the samples and put them on the magnet stand, then remove the supernatant with a P200 pipette.
19. While keeping the tubes on the magnet stand, add 500 μL of Tween-300-wash Buffer to each sample for a wash. Remove the tubes from the magnet stand, shake briefly to mix, put them back onto the magnet stand and wait until clear.
20. Completely remove the liquid, and add 200 μL Tween-300-wash Buffer without mixing for the second wash. Remove the tubes from the magnet stand, shake briefly to mix, put them back onto the magnet stand and wait until clear.
21. Completely remove the liquid, add 300 μL Tagmentation Buffer to each tube and mix well by gently vortexing.
22. Incubate the samples on a heating block at 37°C for 1 hour.

#### Releasing of DNA fragments

23. After the incubation (step 22), add to each tube 10 μL 0.5M EDTA [HUBERLAB # A2937.1000], 3 μL 10% SDS [Roth #CN30.3], and 2.5 μL 20 mg/mL Proteinase K [Thermo Scientific #EO0492] to stop the tagmentation.
24. Mix by vortexing, then incubate at 55°C on a heating block for 1 hour.

#### DNA Extraction with PCI

25. Pre-cool some 100% ethanol [Fisher Chemical E/0665DF/17] and prepare enough 1.5 mL tubes as the number of the samples with 750μL pre-cooled 100% ethanol.
26. To each sample (from step 6.2), add 300 μL phenol:chloroform:isoamyl alcohol 25:24:1 (PCI) [Sigma-Aldrich #77617], and vortex for 2 seconds.
27. Transfer the mixture to a phase-lock tube, centrifuge for 3 minutes at 16,000 x g at room temperature.
28. Add 300 μL chloroform [scharlau #CL02181000] to each phase-lock tube and invert to mix centrifuge for 3 minutes at 16,000 x g at room temperature, to remove the residual phenol in the aqueous phase.
29. Transfer the aqueous layer (the upper layer) to the tubes with 750 μL ethanol (prepared in step 25). Be careful not to stab the pipette tip into the gel of the phase-lock tube.
30. Chill the tubes on ice for 5-10 minutes and centrifuge for 20 minutes at 16,000 x g at 4°C.
31. Gently pour out the supernatant and use a piece of paper towel to completely remove the liquid. The DNA pellet is attached to the wall of the tube although usually not visible.
32. Air dry the tube for 5-10 minutes, then add 1 mL pre-cooled 100% ethanol to wash.
33. Centrifuge at 16,000 x g for 10 minutes at 4°C.
34. Gently pour out the supernatant and use a piece of paper towel to completely remove the liquid. Then air dry the tube until no liquid is visible: it takes about 20 to 30 minutes. Add 22 μL 0.1x TE buffer (1 mM Tris [Roth # AE15.3], pH8, 0.1 mM EDTA [HUBERLAB # A2937.1000]) and vortex well to dissolve the DNA.

#### PCR Amplification for Tagmented Fragments

35. In a PCR tube [Thermo Fisher Scientific #AB0451], mix 21 μL tagmented DNA, 2 μL i5 primer (10 uM) and 2 μL i7 primer (10 uM), and 25 μL NEBNext Hi-Fi 2X PCR mix [NEB #M0541] to make a total final volume of 50 μL. Make sure to use unique combinations of i5 and i7 primers for each library that will be sequence in the same pool. The sequences of i5 and i7 primers are from Kaya-Okur et al. 2020 and can be found in Supplementary Table 2.
36. On a thermocycler, perform the PCR with the program listed in Table 1.

#### PCR purification

37. Once the PCR is completed (step 36), to each sample add 1.1 volume (55 μl) of SPRI beads [NucleoMag # 744970] to each PCR reaction, pipette up and down 10 times to mix well and leave at room temperature for 10 minutes. Note: purification using 1.1 volume of SPRI beads would reduce the amount of short fragments between 200-300bp; Alternatively, one can use 1.3 volume to preserve more fragments, but the short fragments between 200-300 will be dominant in this case
38. Place the PCR tubes on a magnet stand [Invitrogen], wait until the supernatant is clear, then remove the liquid with a 200 μL pipette set at 180 μL.
39. While the tubes stay on the magnet stand, add 180 μL 80% ethanol [Fisher Chemical E/0665DF/17] to wash the beads.
40. Remove the liquid with the 200 μL pipette set at 190 μL, then add another 190 μL of 80% ethanol for the second wash.
41. Remove the liquid with the 200 μL pipette set at 200μL. To remove the liquid attached to the tube wall: briefly spin the tubes, then put them back on the magnet stand, and remove the liquid with the pipette completely.
42. Briefly air dry the beads for less than 5 minutes, then add 25 μL ddH2O to elute the DNA. Vortex and wait for 5 minutes at room temperature.
43. Put the sample back on the magnetic stand, wait until the supernatant becomes clear, then carefully transfer the liquid to a PCR tube [Sarstedt #72.991.002].

#### Library profile and quantitation

44. Use 2 μL of each library to visualize the size distribution with a High Sensitivity D1000 (HSD1000) tape [Agilent 5067-5584] on an Agilent Tapestation 4200. Note: Do not use agrose as the libraries are usually not visible on agarose gels.
45. Select a region from 170 bp to 625 bp to quantify the concentration, as this is the range of size that will be mostly captured by the Illumina sequencers.
46. Proceed to library pooling and next-generation sequencing, or store the libraries at -20°C.

#### Next-generation sequencing

47. Pool the libraries to the desired concentration targeting to reach an equal molarity for each library.
48. Sequence the libraries with a short-read sequencer. The choice of sequencer and flowcell depends on the target read numbers. Aim for 10 – 30 million reads for each H3K27me3 or H3K4me3 or H3K27Ac library.

#### Analysis of CUT&Tag data

Data analysis of CUT&Tag has been performed based on the tutorial from Zheng et al. 2020 [40,41]. Reads were quality-checked using fastqc (0.12.1) and trimmed with fastp (version 0.23.4, -l 30 --trim_poly_x -- poly_x_min_len 32, [42]). Trimmed reads were aligned to the Arabidopsis thaliana reference genome (TAIR10) using bowtie2 (version 2.4.1, options --local --very-sensitive --no-mixed --no-discordant --phred33 -I 10 -X 700, [43]). Duplicate reads were marked with Picard tools (version 3.1.1, broadinstitute.github.io/picard). Fragment count in 500 bp genomic bins are extracted using samtools (version1.17, [44]) and the awk command following the code from Zheng et al. 2020 (awk -F’\t’ ’function abs(x){return ((x < 0.0)-x : x)} {print abs($9)}’ | sort | uniq -c | awk -v OFS=“\t” ’{print $2, $1/2}’, [40,41]) and the correlation indexes were calculated in R (version 4.3.1). Peak calling was done using MACS2 (version 2.2.9.1, options -f BAMPE -q 0.05 --broad --broad-cutoff 0.1 -B -- SPMR, [45]). Peaks from replicates of the same treatment were merged using bedtools (version 2.31.0, [46]). Peaks reproduced in two out of three replicates are considered consensus peaks [47]. To calculate the Fraction of Reads in Peaks (FRiP) score, the .bam files after duplication removal were transformed to .bed files, and the number of reads in peaks was obtained by intersecting the .bed files with peak files using bedtools (version 2.31.0, [46]). The heat plots were plotted using the computeMatrix and plotHeatmap functions from deepTools (version 3.5.3 [48]). The distribution of genomic features on peaks was plotted using ChIPseeker (version 1.38.0, [49]).

**Table 1.**
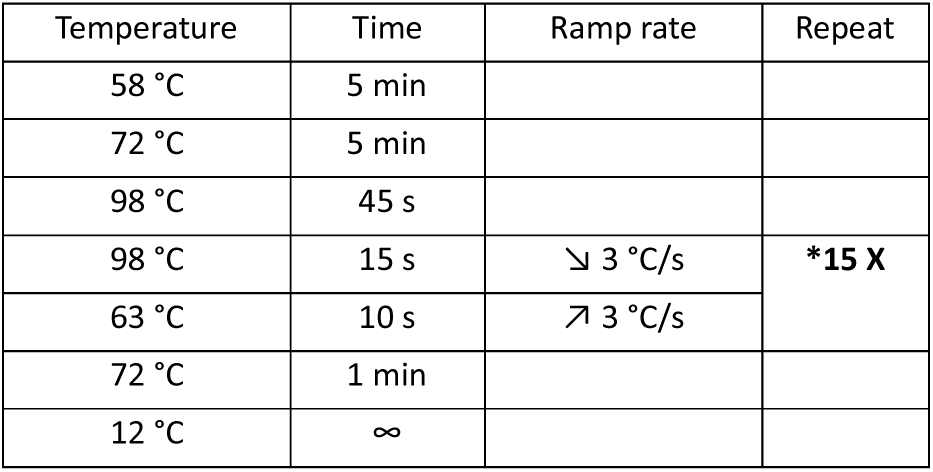
PCR cycle program for CUT&Tag. Note: The cycle number may need optimization depending on the amount of starting material. Please be cautious with increasing the PCR cycle number, as this may lead to unwanted duplication levels that will reduce the quality of your results.

| Temperature | Time | Ramp rate | Repeat |
| --- | --- | --- | --- |
| 58 °C | 5 min |  |  |
| 72 °C | 5 min |  |  |
| 98 °C | 45 s |  |  |
| 98 °C | 15 s | ↘ 3 °C/s | **\*15 X** |
| 63 °C | 10 s | ↗ 3 °C/s |  |
| 72 °C | 1 min |  |  |
| 12 °C | ∞ |  |  |

For visualization on the genome browser, the coverage normalized by sequencing depth was calculated by bamCoverage from deepTools (version 3.5.3, --binSize 10 --normalizeUsing RPGC --effectiveGenomeSize 119481543 --ignoreForNormalization chrC,chrM --extendReads, [48]) and visualized with the Integrative Genomics Viewer (IGV, [50]). To visualize the histone modification with background correction by the H3 controls, .bedGraph files for the histone modifications and control lambda were generated by MACS2 during peak calling with the “-B --SPMR” flag. Background corrected .bedGraph files were generated using the bdgcmp function of MACS2 and converted to .bw files using bedGraphToBigWig (version 445, [51]). The heatmaps were plotted using the computeMatrix and plotHeatmap functions of deepTools (version 3.5.3, [48]).

To associate peaks with genes, peaks are annotated using ChIPseeker (Version 1.38.0, [49]); then, peaks with the “distal intergenic” were excluded from further analysis. The overlaps of marked genes were plotted using VennDiagram (Version 1.7.3, [52]) in R (v4.3.1).

### Comparison of CUT&Tag with ChIP-seq, CUT&RUN, and MNase datasets

To compare our results with previously published ChIP-seq datasets, we downloaded the raw sequencing data from NCBI GEO (GSE84483, Xiao et al., 2017, ChIP-seq data) and ArrayExpress (E-MTAB-10966, Zhu et al., 2023, ChIPmentation data). We processed these public ChIP-seq and ChIPmentation datasets using the same pipeline applied to our CUT&Tag data, with modifications to accommodate single-end sequencing data for GSE84483.

The Venn plots showing the overlap between peaks and the distribution of genomic features on peaks were plotted using ChIPpeakAnno (makeVennDiagram and genomicElementDistribution functions, Version 3.36.1, [53]). Numbers of reads on peaks were counted using featureCounts (Version 2.0.6, [54]). The comparison of peak lengths and read counts in peaks were plotted in R (v4.3.1). The heatmaps around the splicing sites were plotted using the computeMatrix and plotHeatmap functions of deepTools (version 3.5.3, [48]).

The CUT&RUN raw data was downloaded from NCBI GEO (GSE123602) and was processed following the same pipeline applied to our CUT&Tag data. The processed .bigwig files of the MNase-seq data were downloaded from NCBI GEO (GSE205110, [31]).

### Analysis for relations between histone marks and gene expression levels

To get the transcriptome in seedlings of Arabidopsis, we downloaded RNA-seq data from ArrayExpress (E-MTAB-10965, Zhu et al., 2023). Reads were quality-checked using fastqc (0.12.1) and trimmed with fastp (version 0.23.4, -w 16 -l 30 --trim_poly_x --poly_x_min_len 32, [42]).

We used Kallisto (version 0.48.0, [55]) to pseudo-align the trimmed reads to the reference transcriptome. The output of Kallisto, which contains transcripts per million (TPM) numbers, was read and normalized using Sleuth (version 0.30.1, [56]) to obtain normalized transcriptome data. R packages dplyr (version 1.1.4) and ggplot2 (version 3.5.1) were used to process the data and plot transcription levels for genes marked or not marked by different histone modifications.

---

**Supplementary Figure 1.**
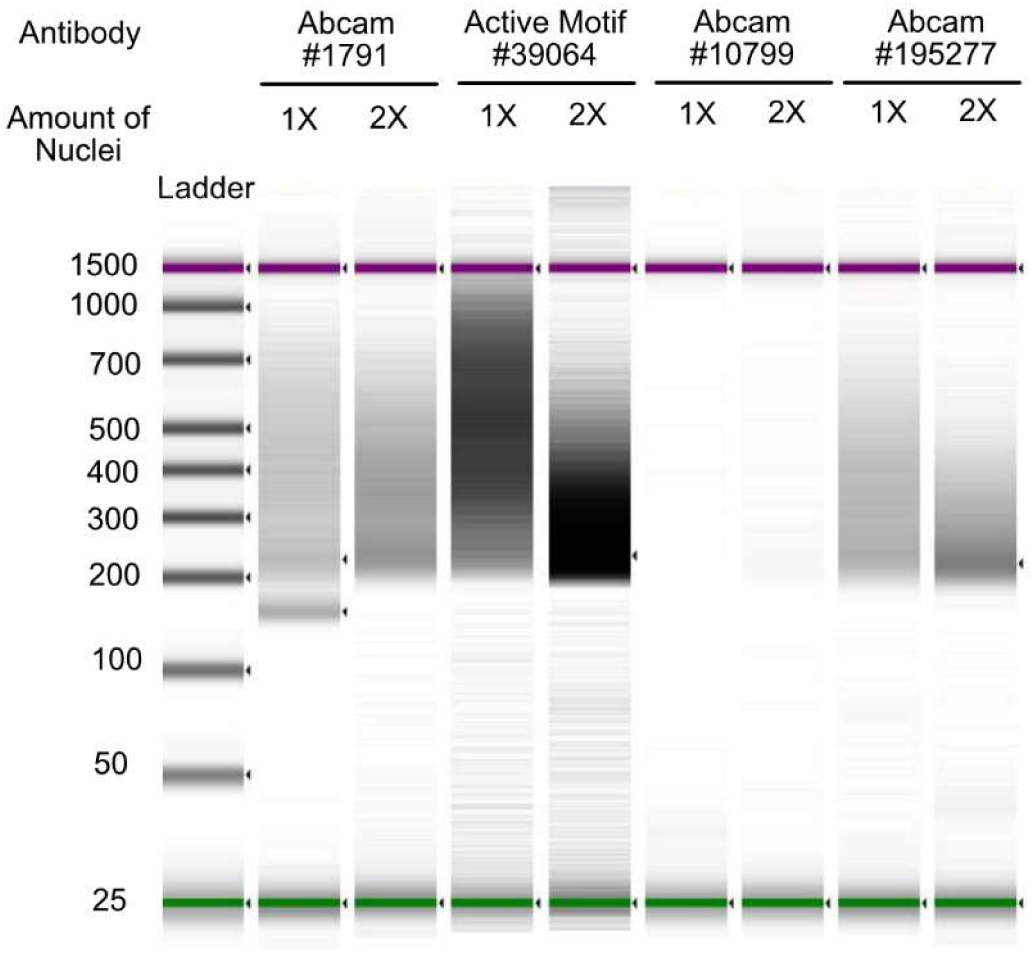
A comparison of the performance of different H3 antibodies in CUT&Tag. Tapestation profiles show the CUT&Tag libraries generated by four different H3 antibodies. Although all the antibodies are ChIP-grade, they show distinctly different performances in CUT&Tag. The antibody Active Motif #39064 generated libraries with much higher concentrations than libraries generated with the other antibodies. One antibody, Abcam #10799, seems to be incompatible with CUT&Tag.

**Supplementary Figure 2.**
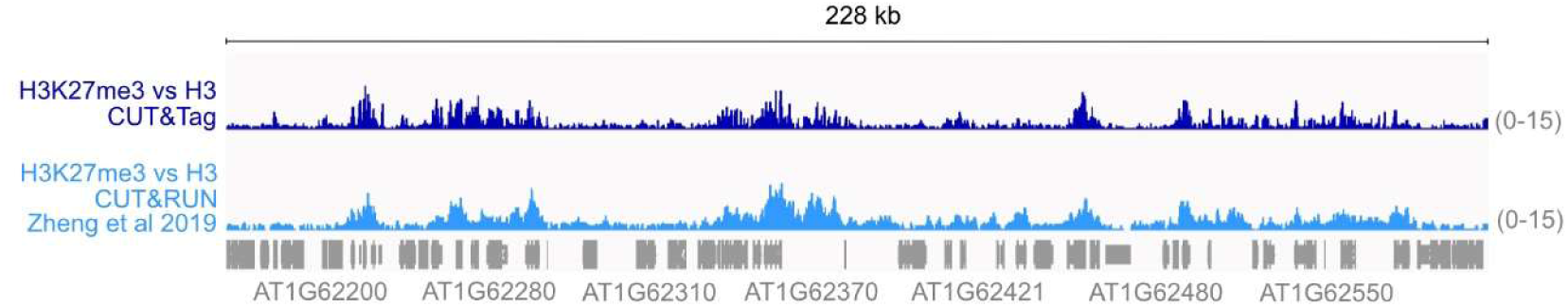
Comparison between CUT&Tag and CUT&RUN H3K27me3 profiles. An overview of a 228 kb genomic window, showing the similarity between the H3K27me3 profile CUT&Tag and that of a previously published CUT&RUN dataset[1].

**Supplementary Figure 3.**
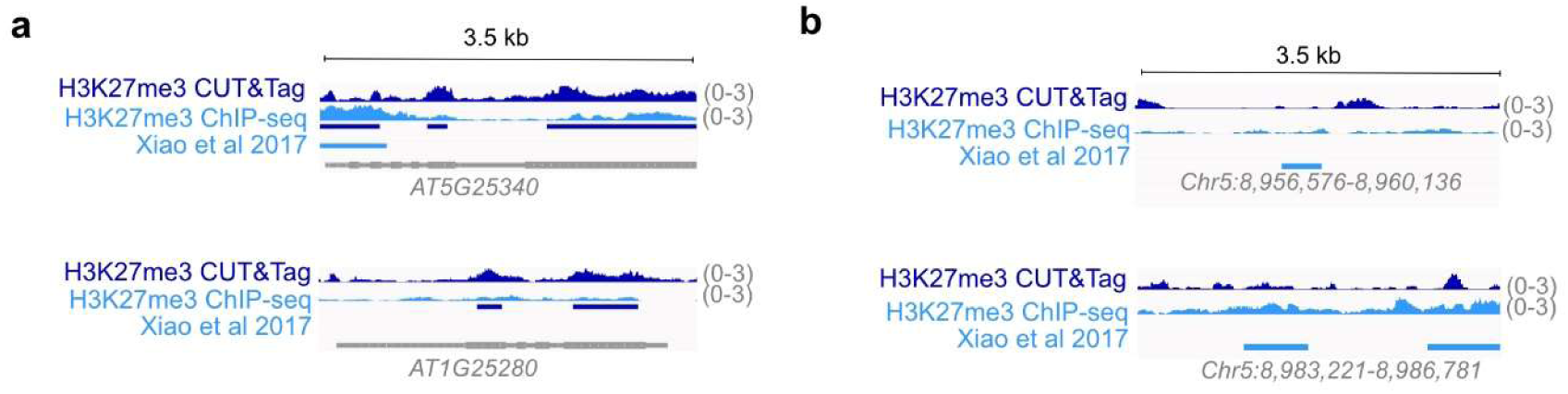
Examples of CUT&Tag-speicific and ChIP-seq-specific peaks. (a) Browser views of two examples of CUT&Tag-specific peaks on exons. (b) Browser views of two examples of intergenic ChIP-specific peaks. Please note that because these peaks are weak, the scales used here are (0-3), which is different from the scales in other figures.

**Supplementary Figure 4.**
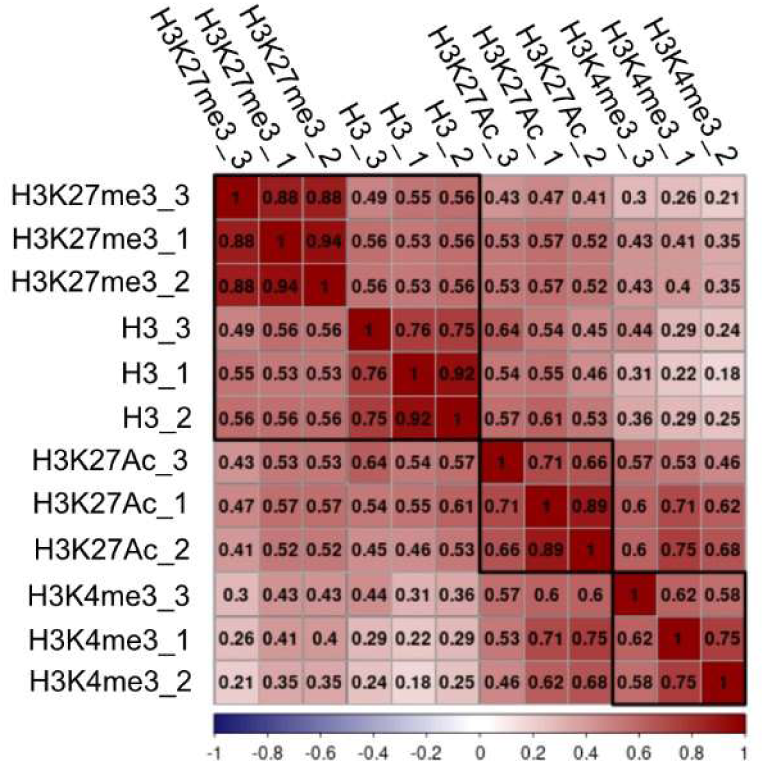
Correlation between CUT&Tag datasets of different histone modifications and the controls. A heat plot shows the correlation between replicates of different histone modifications and the H3 control. Replicates for the same histone modification always cluster together and show high correlations (Pearson correlation coefficient > 0.6), which shows the reproducibility of CUT&Tag experiments.

**Supplementary Figure 5.**
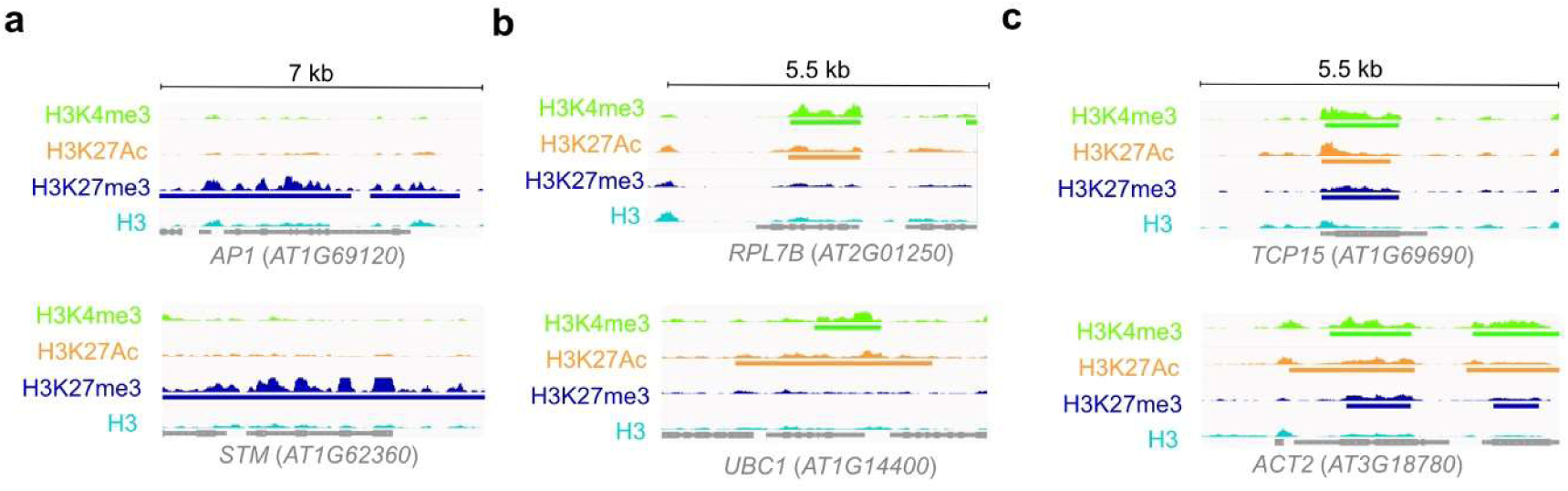
Examples of genes with different chromatin states. Browser views showing examples of genes at different chromatin states: (a) *AP1* (*APETALA1*) and *STM* (*SHOOT MERISTEMLESS*): Polycomb repression; (b) *RPL7B* (*RIBOSOMAL PROTEIN L7B*) and *UBC1* (*UBIQUITIN CARRIER PROTEIN 1*): active transcription; (c) *TCP15* (*TEOSINTE BRANCHED1/CYCLOIDEA/PCF 15*) and *ACT2* (*ACTIN 2*): Mixed.

**Supplementary Figure 6.**
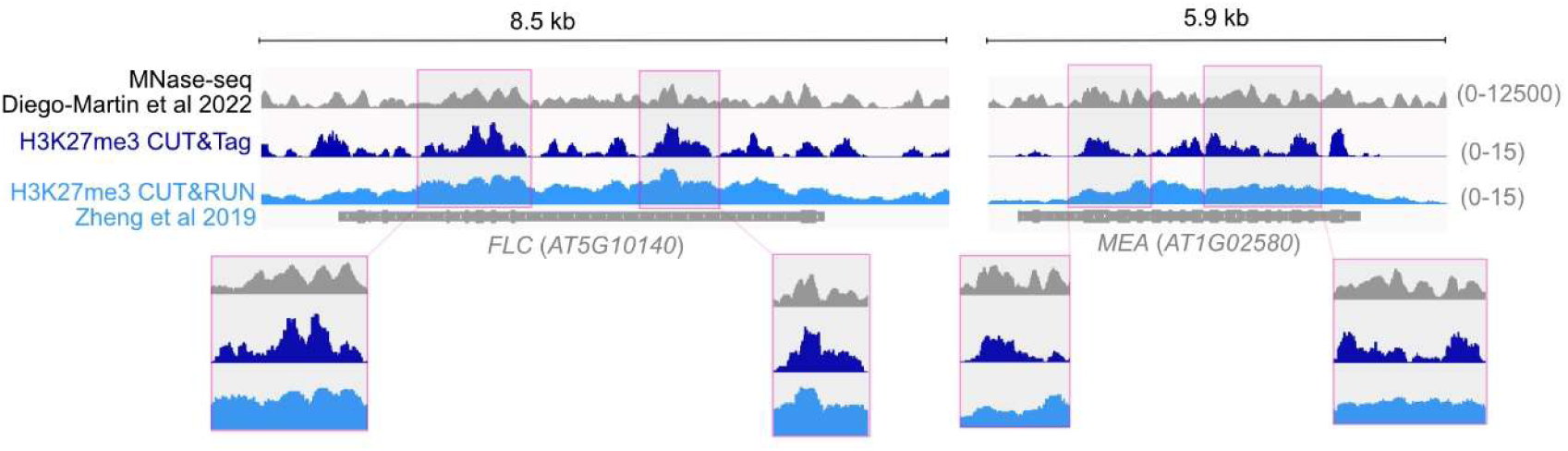
Comparing the resolutions of CUT&Tag and CUT&RUN. Browser views showing H3K27me3 profiles from CUT&Tag and CUT&RUN, together with a nucleosome occupancy profile. The tracks show that the CUT&Tag H3K27me3 profile correlates well with the shape of nucleosome occupancy, while the CUT&RUN H3K27me3 profile correlates with nucleosome occupancy to a weaker degree.

**Supplementary Figure 7.**
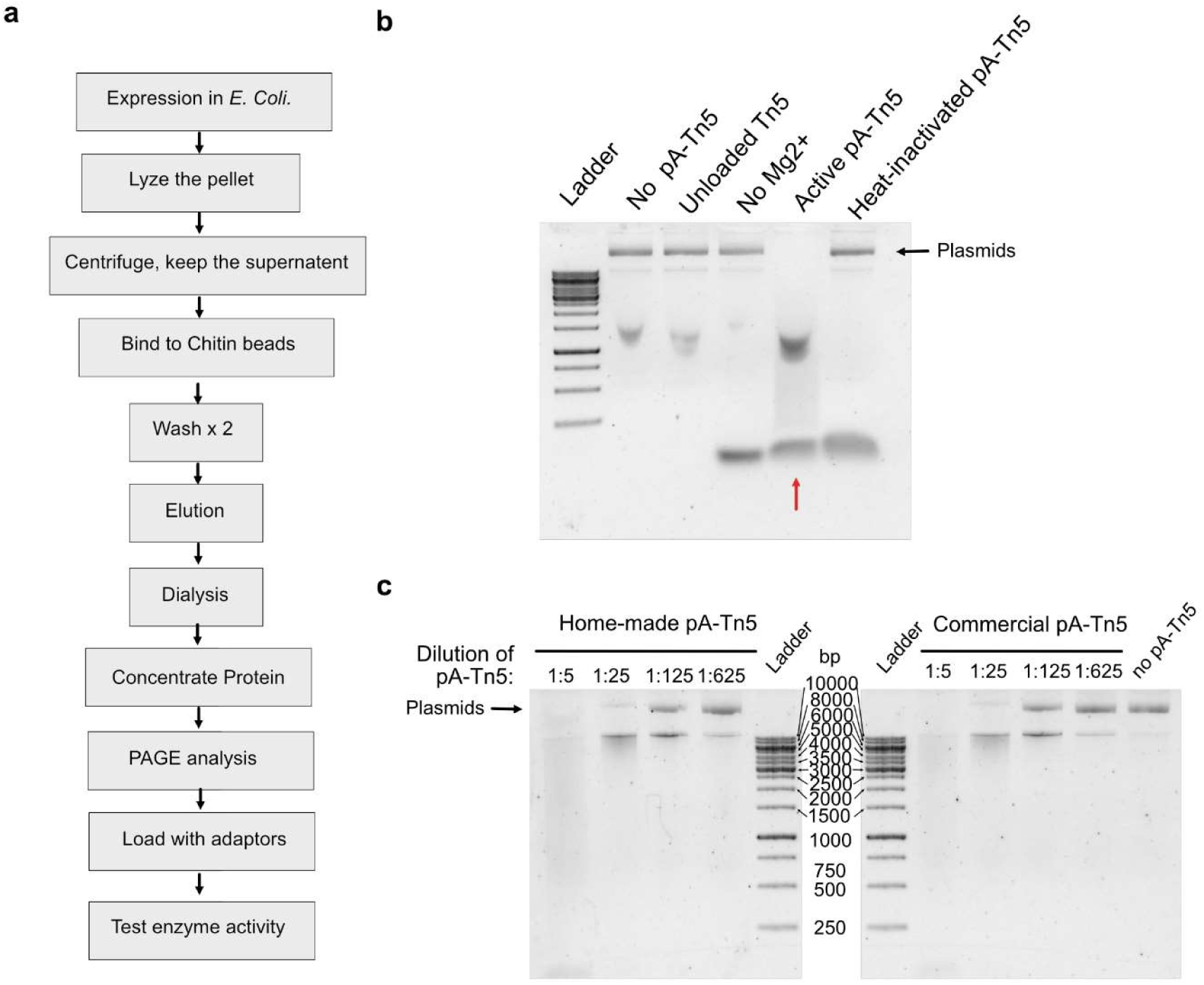
The preparation of pA-Tn5 transposase complex and the tests for its enzymatic activity. (a) The preparation process of pA-Tn5, with the protocol from Li et al. 2021 [2] and Henikoff et al. 2020 [3,4]. (b) Testing the enzyme activity of the loaded pA-Tn5 by digesting a plasmid. As indicated by the red arrow, the plasmid was digested only in the case that pA-Tn5 is loaded, active, and Mg2+ is present. (c) Comparison of home-made pA-Tn5 activity with a commercial pA-Tn5 form Active Motif. The digestion results of the homemade pA-Tn5 are comparable to that of the commercial pA-Tn5, showing their similar activity.

**Supplementary figure 8.**
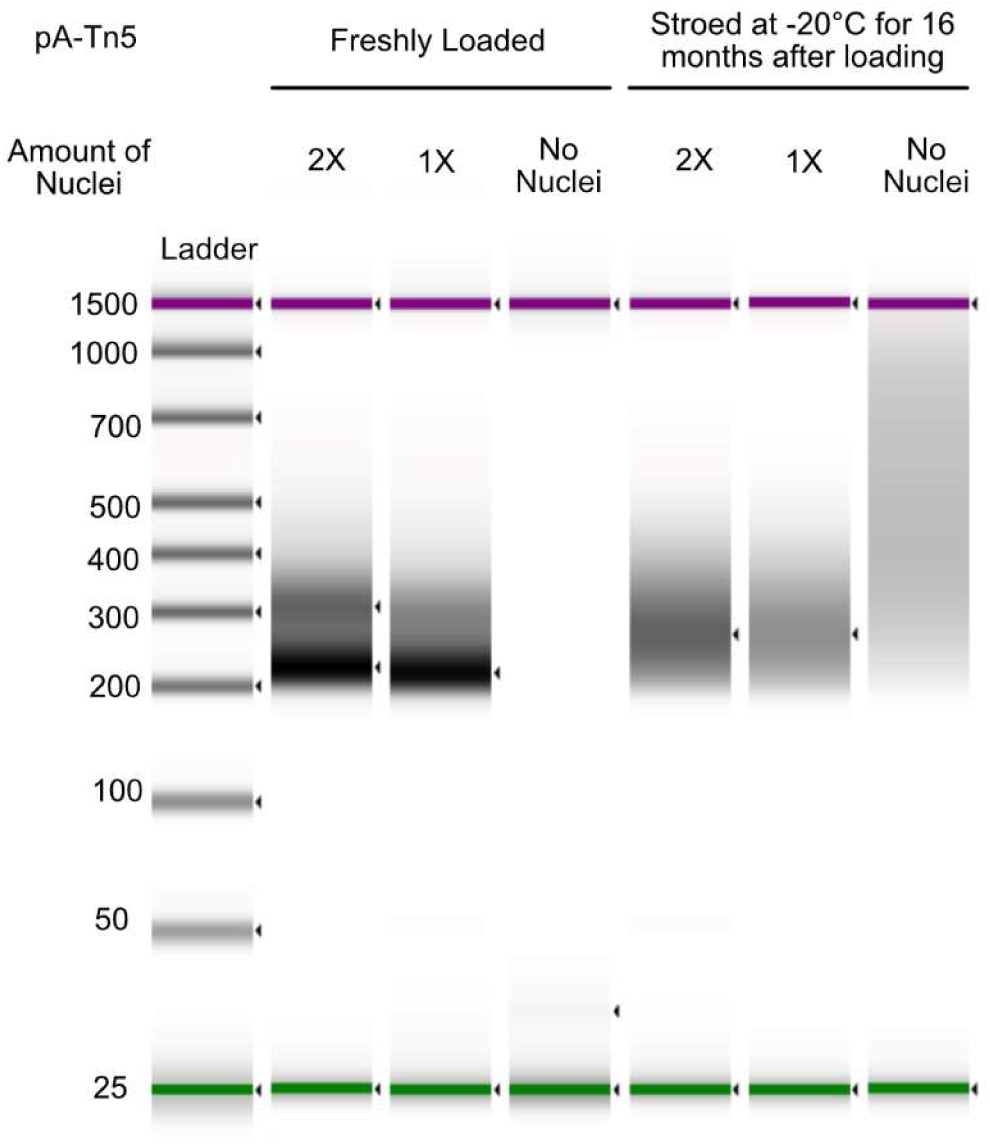
The comparison of performance of freshly loaded pA-Tn5 and pA-Tn5 that has been stored for 16 months after loading. Tapestation profiles show the difference in performance of freshly loaded pA-Tn5 and pA-Tn5 that have been stored for 16 months after loading. The freshly loaded pA-Tn5 generated libraries with nucleosome-ladder patterns from the nuclei of *Arabidopsis* and did not generate a detectable signal when no nuclei were used. The pA-Tn5 that have been stored for 16 months after loading generated signals without nucleosome-ladder patterns and also generated a visible library when no nuclei were used, indicating a high level of tagmented *E. Coli* DNA.

**Supplementary Table 1.** Number of reads after alignment, filter, and deduplication for CUT&Tag datasets.

| Number | Sample | Number of Reads<br>Aligned to TAIR10<br>genome (Million<br>Reads) | Number of Reads<br>passed MAPQ > 10<br>(Miollion Reads) | Number of Reads<br>after Deduplication<br>(Million Reads) |
| --- | --- | --- | --- | --- |
| 323541_09 | H3K27me3_1 | 32.4 | 28.6 | 17.3 |
| 336931_10 | H3K4me3_1 | 49.5 | 33.3 | 14.6 |
| 336931_11 | H3K27Ac_1 | 57.8 | 47.8 | 17.0 |
| 336931_12 | H3_1 | 58.4 | 40.3 | 21.0 |
| 323541_10 | H3K27me3_2 | 33.9 | 30.0 | 18.1 |
| 351531_22 | H3K4me3_2 | 7.4 | 6.6 | 1.9 |
| 351531_23 | H3K27Ac_2 | 32.0 | 29.5 | 10.7 |
| 351531_24 | H3_2 | 42.6 | 35.7 | 22.4 |
| 351531_45 | H3K27me3_3 | 45.9 | 43.3 | 30.5 |
| 351531_34 | H3K4me3_3 | 10.4 | 8.7 | 1.9 |
| 351531_35 | H3K27Ac_3 | 19.2 | 16.5 | 4.9 |
| 351531_36 | H3_3 | 36.1 | 27.8 | 8.7 |

**Supplementary Table 2.**
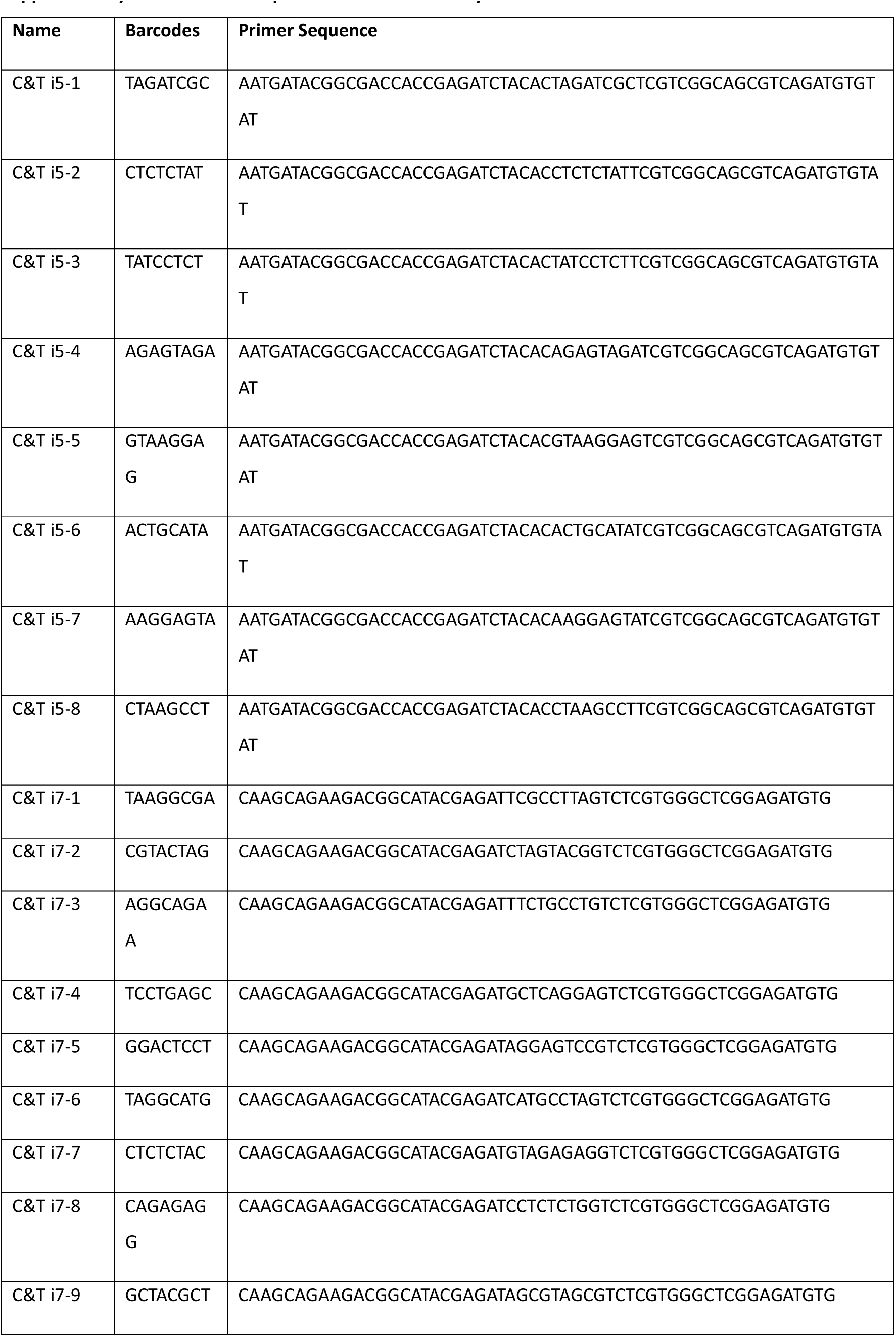
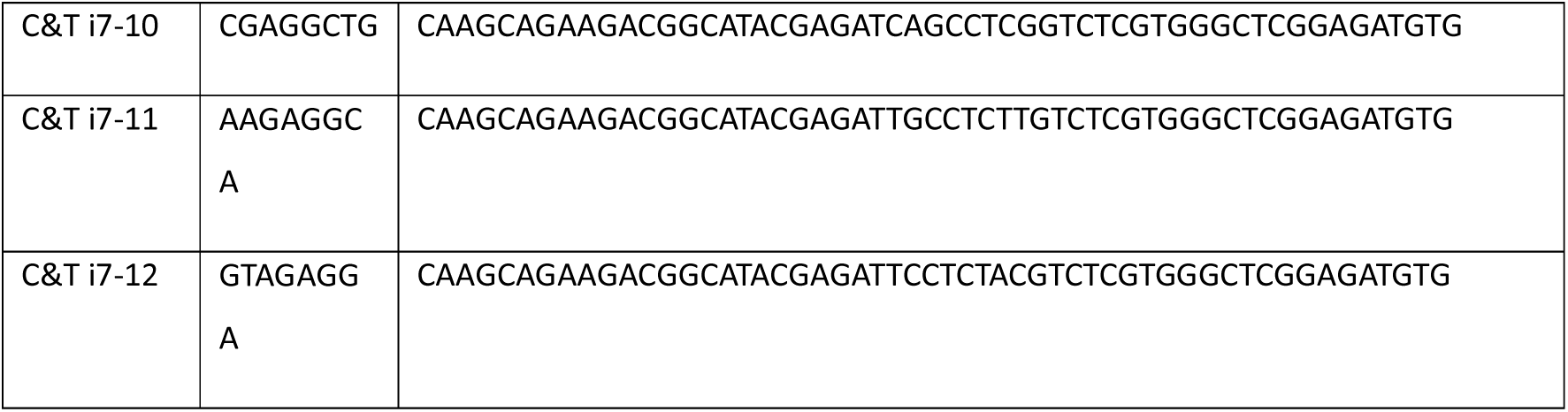
i5 and i7 primers used in this study.

